# Distinguishing classes of neuroactive drugs based on computational physicochemical properties and experimental phenotypic profiling in planarians

**DOI:** 10.1101/2024.07.03.601848

**Authors:** D. Ireland, C. Rabeler, S. Rao, R. J. Richardson, E.-M. S. Collins

**Author notes:** Corresponding authors: (RJR); (E-MSC).

## Abstract

Mental illnesses put a tremendous burden on afflicted individuals and society. Identification of novel drugs to treat such conditions is intrinsically challenging due to the complexity of neuropsychiatric diseases and the need for a systems-level understanding that goes beyond single molecule-target interactions. Thus far, drug discovery approaches focused on target-based *in silico* or *in vitro* high-throughput screening (HTS) have had limited success because they cannot capture pathway interactions or predict how a compound will affect the whole organism. Organismal behavioral testing is needed to fill the gap, but mammalian studies are too time-consuming and cost-prohibitive for the early stages of drug discovery. Behavioral HTS in small organisms promises to address this need and complement *in silico* and *in vitro* HTS to improve the discovery of novel neuroactive compounds. Here, we used cheminformatics and HTS in the freshwater planarian *Dugesia japonica* – an invertebrate system used for neurotoxicant HTS – to evaluate the extent to which complementary insight could be gained from the two data streams. In this pilot study, our goal was to classify 19 neuroactive compounds into their functional categories: antipsychotics, anxiolytics, and antidepressants. Drug classification was performed with the same computational methods, using either physicochemical descriptors or planarian behavioral profiling. As it was not obvious *a priori* which classification method was most suited to this task, we compared the performance of four classification approaches. We used principal coordinate analysis or uniform manifold approximation and projection, each coupled with linear discriminant analysis, and two types of machine learning models –artificial neural net ensembles and support vector machines. Classification based on physicochemical properties had comparable accuracy to classification based on planarian profiling, especially with the machine learning models that all had accuracies of 90-100%. Planarian behavioral HTS correctly identified drugs with multiple therapeutic uses, thus yielding additional information compared to cheminformatics. Given that planarian behavioral HTS is an inexpensive true 3R (refine, reduce, replace) alternative to vertebrate testing and requires zero *a priori* knowledge about a chemical, it is a promising experimental system to complement *in silico* HTS to identify new drug candidates.

**Author summary:** Identifying drugs to treat neuropsychiatric diseases is difficult because the complexity of the human brain remains incompletely understood. Pathway interactions and compensatory mechanisms make it challenging to identify new compounds using computational models and cell-based assays that evaluate potential interactions with specific protein targets. Despite major efforts, neither of these approaches alone nor in combination have been particularly successful in identifying novel neuroactive drugs. Here, we test the hypothesis that rapid behavioral screening using an aquatic invertebrate flatworm, the planarian *Dugesia japonica,* augments the information obtained from computational models based on the physical and chemical properties of neuroactive drugs. Using 19 drugs classified by the vendor as antipsychotics, antidepressants, or anxiolytics, we found that planarian screening could correctly classify most of the drugs based on behavior alone. For compounds known to have multiple therapeutic uses, planarian phenotyping correctly identified the “off-label” class, thereby uncovering effects that were not predicted using the physicochemical properties of the drug alone. This pilot study is the first to show that behavioral phenotyping in a flatworm can be used to classify neuroactive drugs.

## Introduction

Mental illness covers a wide range of conditions that cause significant disturbances in cognition, emotional regulation, or behavior. In 2019, 970 million people or 1 in every 8 people across the world had a mental disorder (1,2). Moreover, the number of people experiencing anxiety or depressive disorders worldwide has significantly increased since the COVID-19 pandemic (3). These statistics are worrisome because psychiatric diseases are intrinsically difficult to treat and pose substantial costs to the affected individuals, families, and society (4,5).

While there is an urgent need for effective therapeutics, neuropsychiatric drug discovery and validation has been stagnant and lags behind that of other diseases (6,7). Computational and *in vitro* methods that try to predict interactions with specific molecular targets are widely used for candidate drug discovery for many illnesses. Although some successful psychiatric drugs, such as fluoxetine, have been identified using this hypothesis-driven approach (8), target-specific methods have been largely insufficient for neuropsychiatric drug discovery due to the complex and still enigmatic pathologies of these disorders (6,9,10). Mental illnesses, such as schizophrenia and major depressive disorder, can stem from polygenic and non-genetic etiologies that likely depend on an interplay between many different molecular targets (reviewed in (11)). Consequently, successful psychiatric drugs tend to be neuroactive compounds with multiple pharmacological targets (12). Thus, organismal models that provide systems-level insight into neuronal function are needed. However, mammalian tests are prohibitively time and cost intensive to screen the large number of chemicals evaluated during the early stages of lead identification (10,13).

One possible solution to integrate organismal information into first-tier lead identification is to use behavioral phenotyping with small organisms. Fish larvae, worms, and flies are inexpensive to maintain compared to mammals and lend themselves to high-throughput screening (HTS) (reviewed in (14,15)). Because phenotypic drug discovery does not rely on mechanistic knowledge and can identify polypharmacological drugs (16,17), it is especially well-suited for neuropsychiatric drug discovery.

Behavioral barcodes, which reduce the multidimensional phenotypic information into a string of numeric features, provide a quantitative readout of a phenotype (18). Using statistical tools such as hierarchical clustering or multidimensional scaling, behavioral profiling can be used to identify patterns characterizing drug classes and have been used to predict the effects of novel chemicals for psychiatric uses (17–19). For example, behavioral barcoding derived from a series of motor responses of zebrafish larvae to different auditory and visual stimuli was used to identify potential novel antipsychotic compounds by comparing to the phenotypic profile produced by classical antipsychotics, such as haloperidol (20). Similar methodology has also been employed to identify novel monoamine oxidase and acetylcholinesterase inhibitors (21) and sedatives that induce paradoxical excitation (22). However, the zebrafish larvae used for these studies were > 7 days old and thus qualify as animal tests that require Institutional Animal Care & Use Committee approval (23).

Here, we test the hypothesis that HTS using an aquatic invertebrate, the planarian *Dugesia japonica,* allows for unbiased classification of neuroactive drugs and augments the information obtained from the physical and chemical properties of the drugs. Freshwater planarians are small (a few mm long) flatworms, have a long history of use in neurotoxicity and neuropharmacology studies (reviewed e.g., in (24–27)), and show promise as a tool for neuroactive drug discovery. The planarian nervous system shares many of the same neurotransmitters and cell types as the mammalian nervous system, yet remains tractable, consisting of approximately 10,000 neurons (26,28), making the planarian nervous system of intermediate size and complexity compared to nematodes and zebrafish. Genomes and transcriptomes are publicly available (29,30), and RNA interference can be used to connect genotypes and phenotypes (31–38). Planarians display stereotypical behavioral responses to drugs of abuse and prescription drugs (39–41). The development of computational methods for quantitative assessment of these behaviors (42–44) have reignited interest in planarian behavioral studies (15,26–28,45). We have shown that the asexual planarian *D. japonica* is well-suited for behavioral HTS (43,44,46,47) and that the phenotypic profiles gained from planarian HTS can distinguish between different chemical classes/modes of action for neurotoxic compounds (44,46,48). Planarian HTS has unique strengths that complement HTS using zebrafish larvae or roundworms for developmental neurotoxicology (14). Adult nervous system function can be evaluated in planarians at a fraction of the time and cost required for zebrafish.

We hypothesized that *D. japonica* behavioral phenotyping would be a good model for the identification of prospective neuroactive drugs. To test this, we used our robotic screening platform to determine how well our battery of phenotypic endpoints could distinguish between classes of known neuropsychiatric drugs and add value to a classification of these compounds based on their physicochemical properties. We studied 19 neuroactive compounds from three functional classes: antipsychotics, anxiolytics, and antidepressants. We classified these compounds using either physicochemical descriptors or planarian behavioral profiling using four different classification approaches. We used principal coordinate analysis (PCoA) or uniform manifold approximation and projection (UMAP), each coupled with linear discriminant analysis (LDA), and two types of machine learning models - artificial neural net ensembles (ANNE) and support vector machines (SVMs). We found that the classification based on physicochemical properties had comparable accuracy to planarian profiling. Planarian profiling correctly identified polytherapeutic drugs and thus yielded additional information that would have been missed by cheminformatics. Thus, combining these two approaches may be an economical and useful method for identifying novel drug candidates.

## Results and Discussion

The purpose of this study was to investigate the extent to which members of neuroactive drug classes can be identified and distinguished using computational classification using either cheminformatics or *in vivo* behavioral phenotyping in planarians. To do this, we used a set of 19 neuroactive drugs that were functionally defined by the supplier as one of three categories: antidepressants (7), antipsychotics (7), or anxiolytics (5), (Table 1, Fig 1).

**Fig 1.**
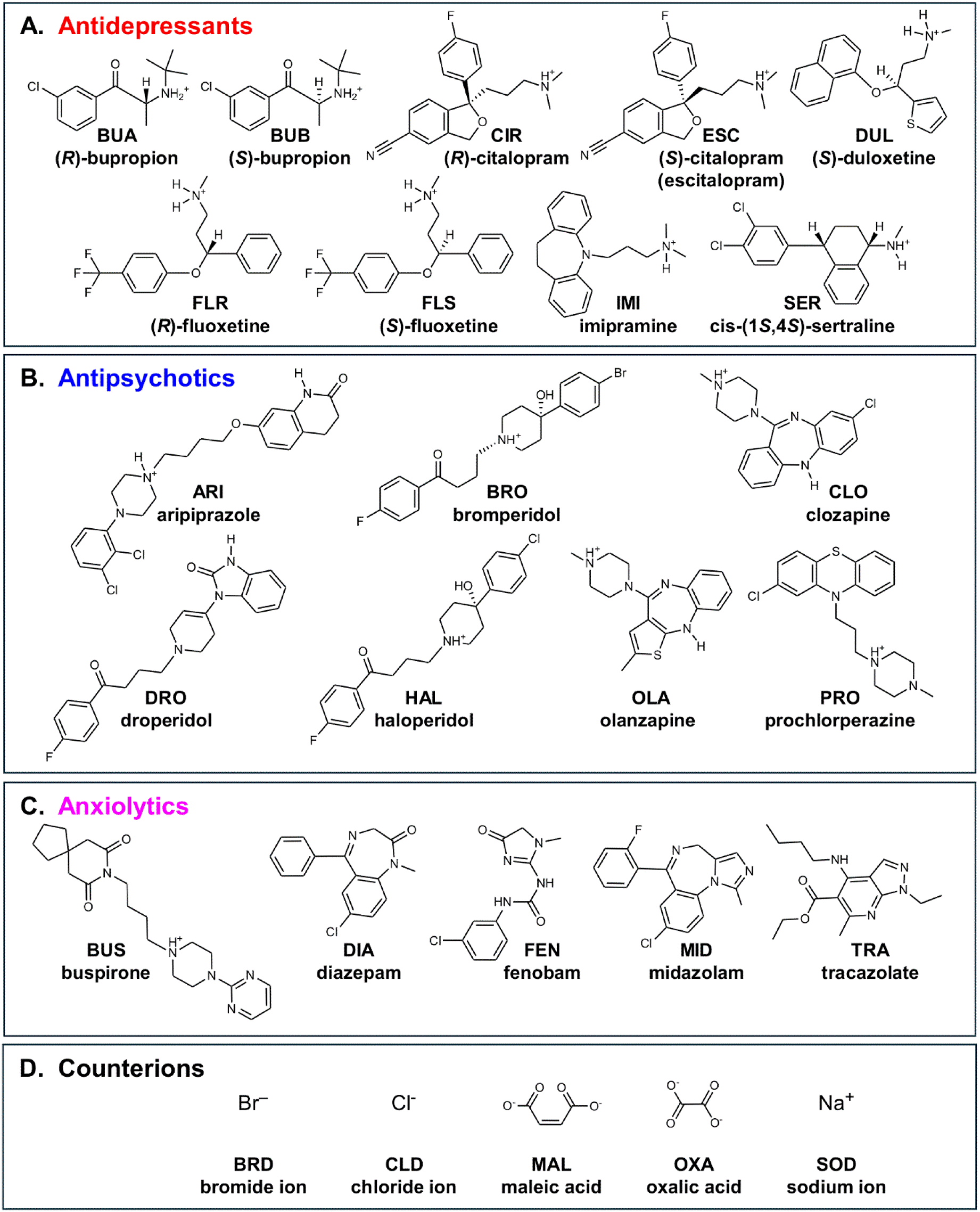
Chemical Structures. Molecular structures depicted as 2D with stereochemistry indicated by wedged bonds are provided for the different chemical classes: A) Antidepressants, B) Antipsychotics, C) Anxiolytics, and D) Counterions. Counterions are not shown with their respective drug. Protonation states were based on the dominant form at pH 7.4 as determined by the protonation module in the ChemAxon Marvin suite (https://www.chemaxon.com).

**Table 1.** Study compounds.

| CAS # | ID | Common Name<sup>a</sup> | Class<sup>b</sup> | Source<sup>c</sup> | Purity (Method)<sup>d</sup> | Solvent<sup>e</sup> | Study Type<sup>f</sup> | Tested Concentrations (units) |
| --- | --- | --- | --- | --- | --- | --- | --- | --- |
| 129722-12-9 | ARI | Aripiprazole·HCl | 2 | A | ≥98% (HPLC) | DMSO | E,C | 1, 3.16, 10, 31.6, 100 (μM) |
| 10457-90-6 | BRO | Bromperidol | 2 | A | 100% (EP reference standard) | DMSO | E,C | 0.316, 1, 3.16, 10, 31.6, 100 (μM) |
| 437723-96-1 | BUA | (R)-isomer of BUP | 1 | NA | NA | NA | C | NA |
| 324548-43-8 | BUB | (S)-isomer of BUP | 1 | NA | NA | NA | C | NA |
| 31677-93-7 | BUP | Bupropion·HCl<br>Racemic mixture | 1 | A | ≥98% (HPLC) | water | E | 10, 31.6, 100, 316, 1000 (μM) |
| 33386-08-2 | BUS | Buspirone·HCl | 3 | A | ≥99% (TLC) | water | E,C | 0.316, 1, 3.16, 10, 31.6, 100 (μM) |
| 59729-32-7 | CIT | Citalopram·HBr<br>Racemic mixture | 1 | A | quality level<br>300; certified<br>reference<br>material<br>(HPLC) | DMSO | E | 1, 3.16, 10, 31.6,<br>100, 316, 1000<br>(µM) |
| 128196-02-1 | CIR | (R)-isomer of CIT | 1 | NA | NA | NA | C | NA |
| 5786-21-0 | CLO | Clozapine | 2 | B | ≥99%<br>(HPLC) | DMSO | E,C | 0.1, 0.316, 1,<br>3.16, 10, 31.6,<br>100 (µM) |
| 439-14-5 | DIA | Diazepam | 3 | C | 99.3%<br>(GC) | DMSO | E,C | 1, 3.16, 10, 31.6,<br>100 (µM) |
| 548-73-2 | DRO | Droperidol | 2 | A | ≥98%<br>(TLC) | DMSO | E,C | 0.1, 0.316, 1,<br>3.16, 10, 31.6,<br>100 (µM) |
| 136434-34-9 | DUL | Duloxetine·HCl<br>(S)-isomer | 1 | A | ≥98%<br>(HPLC) | DMSO | E,C | 1, 3.16, 10, 31.6,<br>100 (µM) |
| 219861-08-2 | ESC | Escitalopram·oxalate<br>(S)-isomer of CIT | 1 | A | ≥98%<br>(HPLC) | DMSO | E,C | 1, 3.16, 10, 31.6,<br>100, 316, 1000<br>(µM) |
| 57653-26-6 | FEN | Fenobam | 3 | A | ≥98%<br>(HPLC) | DMSO | E,C | 56.2, 100, 178,<br>316, 562 (µM) |
| 100568-03-4 | FLR | (R)-isomer of FLU | 1 | NA | NA | NA | C | NA |
| 100568-02-3 | FLS | (S)-isomer of FLU | 1 | NA | NA | NA | C | NA |
| 56296-78-7 | FLU | Fluoxetine·HCl<br>Racemic mixture | 1 | A | ≥98%<br>(TLC) | DMSO | E,C | 1, 3.16, 10, 31.6,<br>100 (µM) |
| 52-86-8 | HAL | Haloperidol | 2 | A | ≥98%<br>(TLC) | DMSO | E,C | 1, 3.16, 10, 31.6,<br>100 (µM) |
| 113-52-0 | IMI | Imipramine·HCl | 1 | A | ≥99%<br>(TLC) | DMSO | E,C | 0.316, 1, 3.16, 10,<br>31.6 (µM) |
| 110-16-7 | MAL | Maleic acid | 4 | A | ≥99%<br>(HPLC) | DMSO | E,C | 200 (µM) |
| 59467-70-8 | MID | Midazolam | 3 | A | 100%<br>(USP<br>reference<br>standard) | DMSO | E,C | 10, 17.8, 31.6,<br>56.2, 100 (µM) |
| 132539-06-1 | OLA | Olanzapine | 2 | A | ≥98%<br>(HPLC) | DMSO | E,C | 0.001, 0.00316,<br>0.01, 0.0316, 0.1,<br>0.316, 1, 3.16, 10,<br>31.6, 100, 316<br>(µM) |
| 144-62-7 | OXA | Oxalic acid | 4 | A | Analytical<br>standard<br>(redox<br>titration) | DMSO | E,C | 100, 316, 1000<br>(µM) |
| 84-02-6 | PRO | Prochlorperazine·<br>dimaleate | 2 | A | ≥98%<br>(TLC) | DMSO | E,C | 0.316, 1, 3.16, 10,<br>31.6, 100 (µM) |
| 79559-97-0 | SER | Sertraline·HCl<br>*cis*-(1*S*,4*S*)-isomer | 1 | A | ≥98%<br>(HPLC) | DMSO | E,C | 1, 3.16, 10, 31.6,<br>100 (µM) |
| 7647-15-6 | SOB | Sodium bromide | 4 | A | ≥99.9%<br>(trace metal<br>analysis) | DMSO | E | 100, 316, 1000<br>(µM) |
| 7647-14-5 | SOC | Sodium chloride | 4 | A | ≥99.0%<br>(titration with<br>AgNO<sub>3</sub>) | water | E | 3.47 (mM) |
| 1135210-68-2 | TRA | Tracazolate·HCl | 3 | B | ≥99.0%<br>(HPLC) | DMSO | E,C | 1, 3.16, 10, 31.6,<br>100 (μM) |

<sup>a</sup> All common names are those provided in the National Institutes of Health Global Substance Registration System. Except for maleic acid, oxalic acid, sodium bromide and D-sorbitol, all names also represent the recommended International Non-Proprietary Name.
<sup>b</sup> Primary functional pharmacological class as indicated by the manufacturer and literature: 1 = antidepressant; 2 = antipsychotic; 3 = anxiolytic; 4 = counterion control.
<sup>c</sup> Source of chemicals used in experimental determinations: A = Sigma-Aldrich; B = Tocris; C = Spectrum Chemicals; NA = not applicable (molecular structures used only for *in silico* determinations).
<sup>d</sup> Method used by the supplier to assess purity. HPLC: High-performance liquid chromatography; EP: European Pharmacopeia; GC: Gas chromatography; TLC: Thin layer chromatography; USP: US Pharmacopeia.
<sup>e</sup> Solvent used for stock solutions.
<sup>f</sup> Study type: E = experimental; C = computational. For computational studies, counterions were not present.

We assigned each chemical a barcode consisting of a numerical string based on either their computed physicochemical properties or responses observed in planarian phenotypic screening. We then used the same computational pipeline to study how well these compounds could be accurately classified based on either *in silico* cheminformatics or *in vivo* planarian behavioral phenotyping. We compared the results obtained from four different computational methods, including PCoA followed by LDA (PCoA-LDA), UMAP followed by LDA (UMAP-LDA), and two machine-learning approaches (ANNE and SVMs) (Fig 2), to investigate whether a specific method would work best on either or both types of data. In the Methods section, we provide background on each method and explain why we chose to apply it to our data.

**Fig 2.**
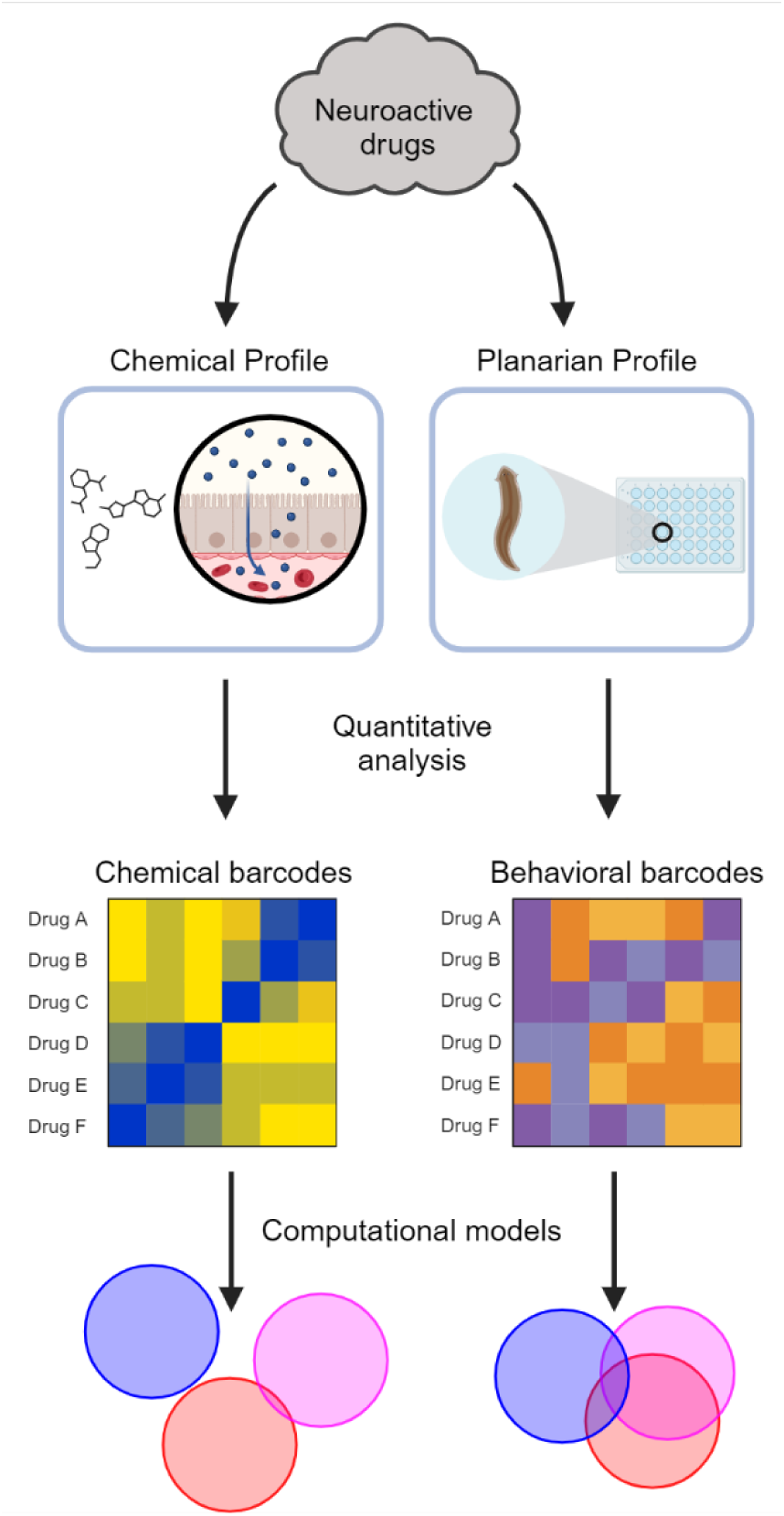
Study overview. Chemical and behavioral profiles were determined for 19 neuroactive drugs, consisting of 7 antidepressants, 7 antipsychotics, and 5 anxiolytics. Using quantitative analyses, we then determined molecular and phenotypic barcodes for each compound. These barcodes were used in the same computational models to determine how well each method performed at classifying the 3 neuroactive drug classes. Created with BioRender.com.

### Classification based on physicochemical properties

The neuroactive drugs studied here represent a diverse range of chemical structures, as shown by their differing Tanimoto similarity coefficients, compared to ARI as a reference compound (Fig 3). While some clustering by functional pharmacological class is visible, several compounds (e.g., the antipsychotics OLA and CLO and most anxiolytics) are structurally distinct from the other members of their class. This structural diversity reflects the purpose of this library – to serve as a pilot to determine the extent to which our models could assign the compounds to their functional pharmacological classes despite structural dissimilarities within each class.

**Fig 3.**
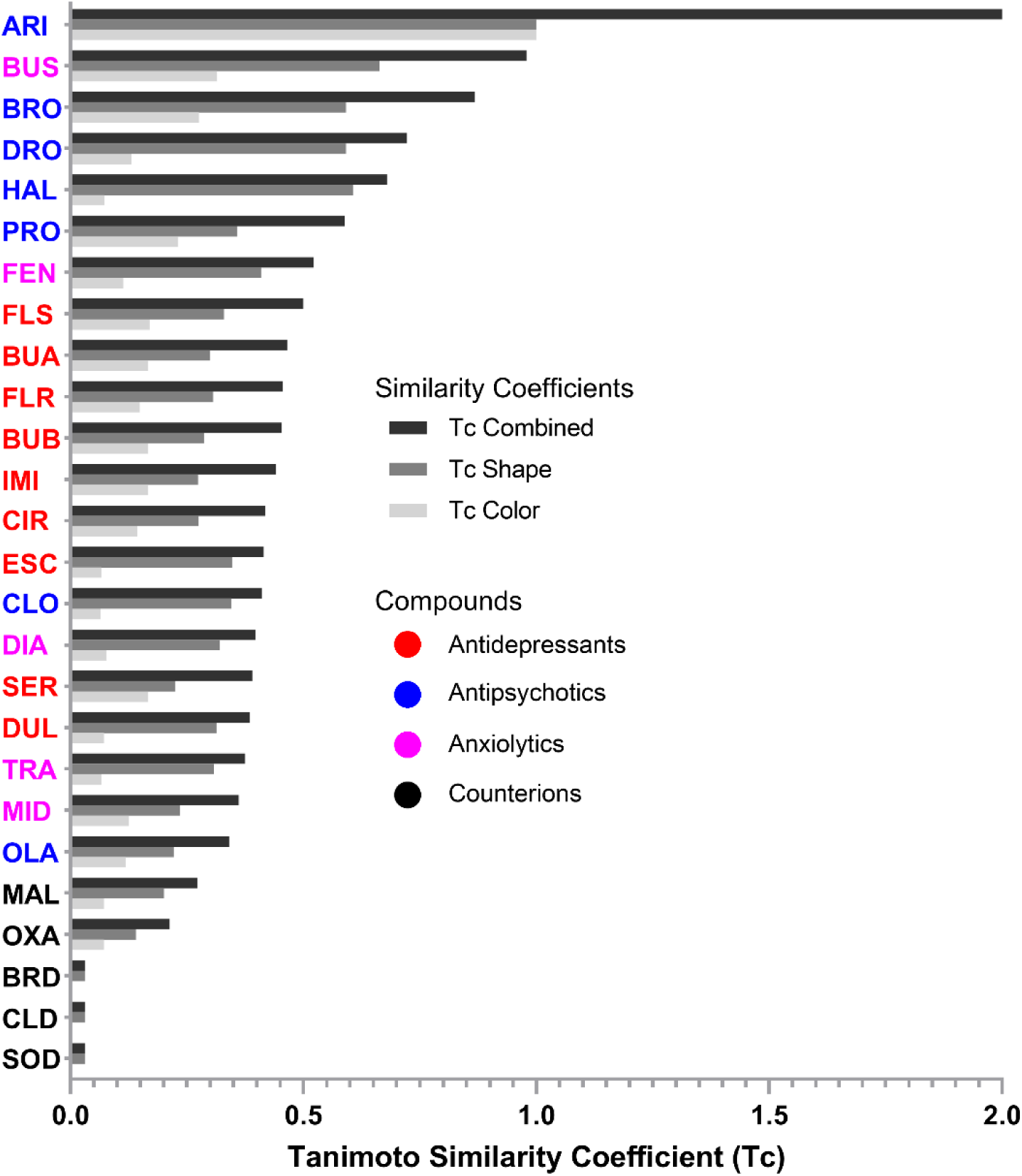
Tanimoto similarity coefficients. Chemical similarity was quantified using the Tanimoto coefficient (Tc). Overlaps were determined after 3D alignment of each structure with the reference compound, ARI. Tc Shape refers to similarity based on overlap of molecular volume. Tc Color refers to similarity based on overlap of 6 pharmacophores (H-bond acceptor, H-bond donor, anion, cation, hydrophobe, or ring). Tc Combined = Tc Shape + Tc Color. Range of Tc Shape and Tc Color = (0,1); range of Tc Combined = (0,2). Chemicals are ordered by their Tc Combined scores and color-coded by functional class.

As shown in Table 1, some drugs (BUP, CIT, FLU) are mixtures of stereoisomers. Thus, we had to consider whether to treat these chemicals as 2D structures, without considering stereochemistry, or as 3D structures, considering each stereoisomer separately thereby increasing the number of entities considered. Because the number of compounds affects the classification and we wanted to keep the data size comparable to the planarian data, which considers each tested compound as a single entity, we present the classifications based on 2D structural features here without consideration of stereochemistry (Figs 4-6). Results from using 3D chemical features, considering each stereoisomer separately, are shown in S1-3 Figs. The 3D models show similar, though slightly better, classification accuracies than the models using only 2D features, which may be due to the increased information per compound and/or the increase in the number of compounds (18 versus 21) compared to 2D. Of note, when considering 2D structures, CIR and ESC were treated as one structure, which we labeled CIT, leading to only 18 drugs in these models.

**Fig 4.**
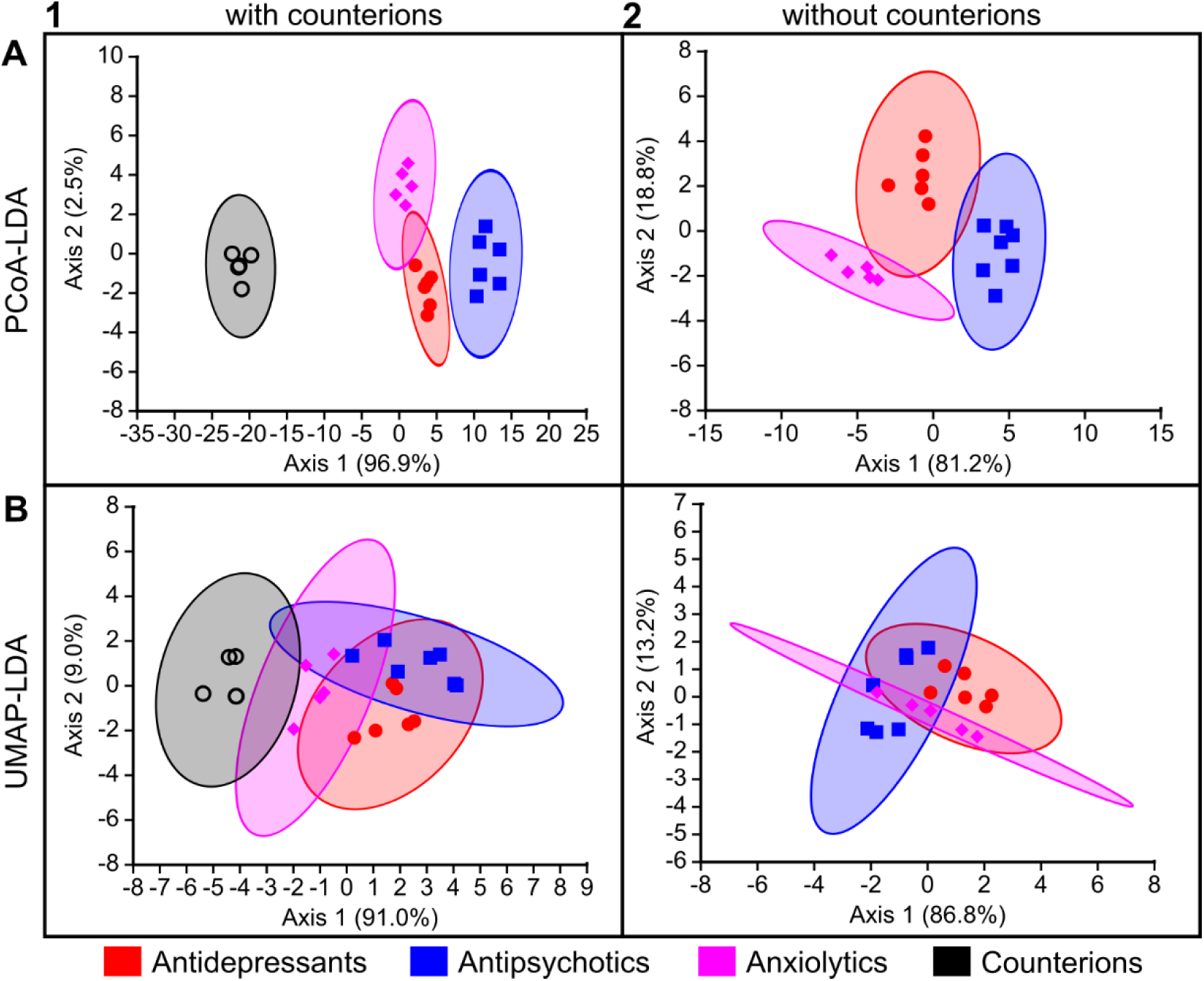
Functional pharmacological classification of the study compounds based on their 2D chemical descriptors using LDA-based approaches. Computational methods are arranged by rows: (A) PCoA-LDA; (B) UMAP-LDA. Counterion inclusion/exclusion is arranged by columns: (1) with counterions; (2) without counterions. Ellipses represent 95% confidence intervals. The axes show the percentage of the total eigenvalues; this does not sum to 100% in (A1) because there was a third axis that accounted for the remaining 0.60%.

Comparisons were made both with and without the counterions to determine how inclusion of these presumedly null-effect compounds would affect neuroactive drug classification. ANNE and SVMs were each initially run across 10 different models (S1-9 Tables) and then the best performing model for each was chosen (see Methods) for comparison across methods.

The accuracy of these models on the small data set used here was high and comparable between PCoA-LDA, ANNE, and SVMs (Figs 5-6). Overall, the UMAP-LDA performed worse than the other three models on all data sets (2D, 3D +/- counterions). Interestingly, when using the 2D chemical features and excluding the counterions, UMAP-LDA misclassified most anxiolytics (Fig 6B). Only FEN was correctly classified. Generally, classification accuracies were improved by including the counterions, which may be due to the increased number of chemicals. When comparing the misclassifications from the 2D and 3D chemical descriptors, we found that the antidepressant BUP was the most frequently misclassified drug across the different methods when using 2D structural features (Figs 5-6).

**Fig 5.**
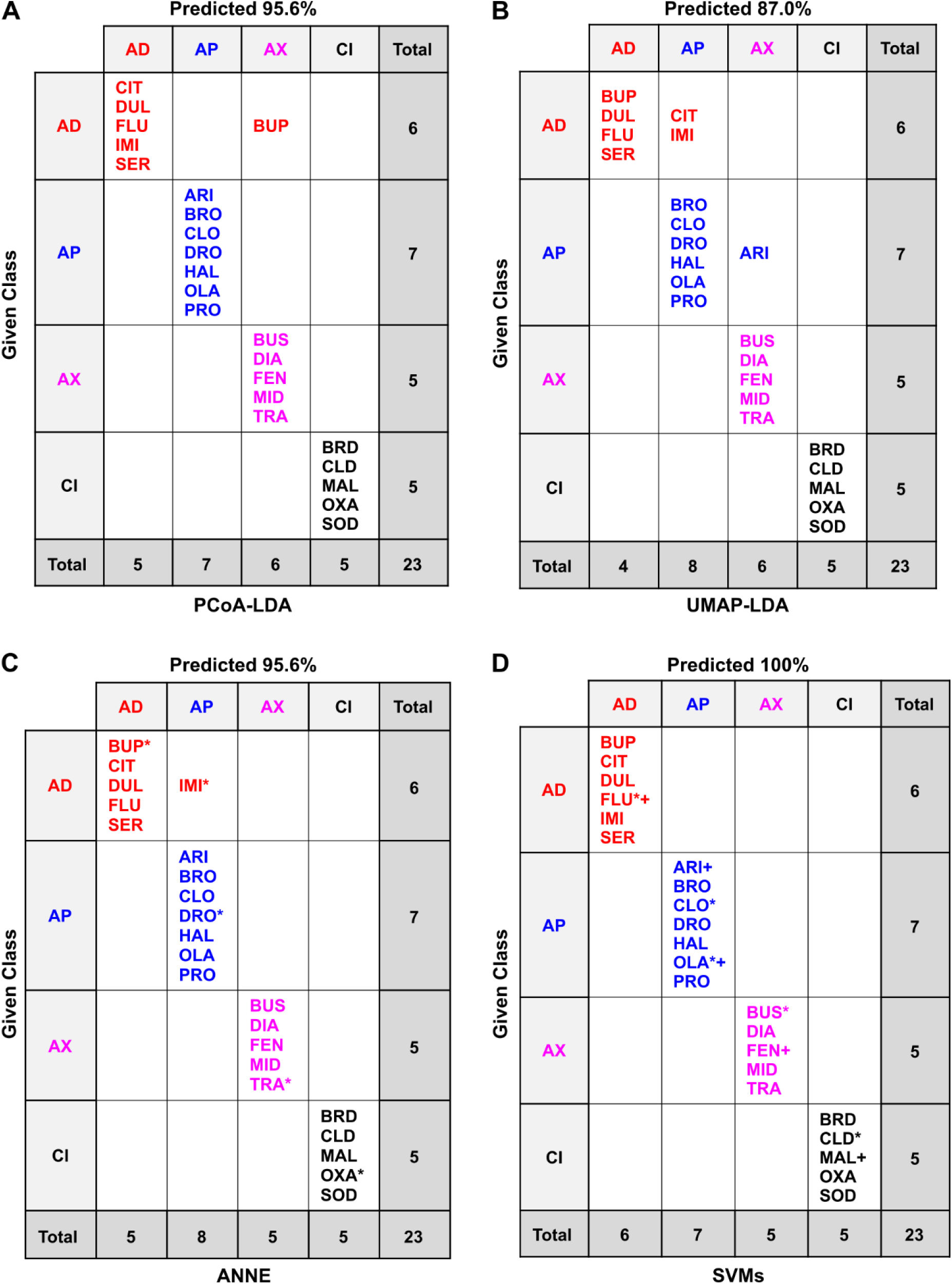
Confusion matrices of the classification methods based on 2D chemical descriptors of the drugs and counterions. Confusion matrices for the different classification methods: (A) PCoA-LDA; (B) UMAP-LDA; (C) ANNE; (D) SVMs; AD: antidepressant, AP: antipsychotic, AC: anxiolytic; CI: counterion. In A and B, predicted accuracy was calculated following an exhaustive jackknifing. In C and D, predicted accuracy refers to the overall accuracy of the best of 10 models of each type. *indicates randomly chosen members of the test set. In (D) two models were tied for the best so that the test set members are marked with both * (model 07_2i) and + (model 09_2i). See the Methods for model selection. Test set accuracies were 80.0% for ANNE (S2 Table) and 100% for SVMs (S4 Table).

**Fig 6.**
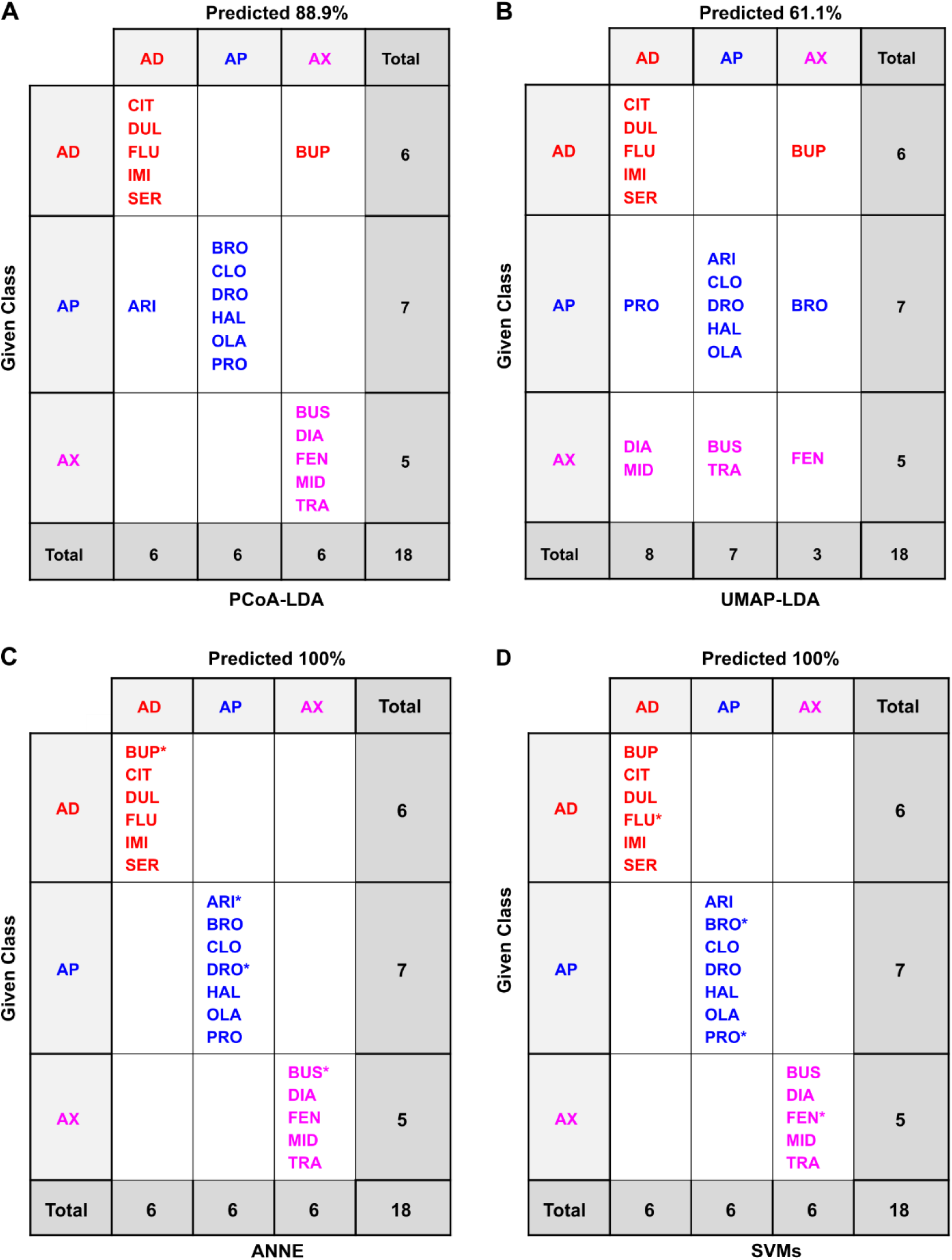
Confusion matrices of the classification methods based on 2D chemical descriptors of the drugs only. Confusion matrices for the different classification methods: (A) PCoA-LDA; (B) UMAP-LDA; (C) ANNE; (D) SVMs; AD: antidepressant, AP: antipsychotic, AC: anxiolytic. In A and B, predicted accuracy was calculated following an exhaustive jackknifing. In C and D, predicated accuracy refers to the overall accuracy of the best of 10 models of each type. *indicates randomly chosen members of the test set. See the Methods for model selection. Test set accuracies were 100% for ANNE (S3 Table) and 100% for SVMs (S5 Table).

In contrast, the classifications based on 3D chemical structures did not misclassify any antidepressants, including the BUP stereoisomers (BUA and BUB), and most frequently misclassified anxiolytics (S2 and 3 Figs). When considering 3D chemical features, the most misclassified chemical across the different methods was the anxiolytic BUS (S2-3 Figs). BUS has a very different structure compared to the other anxiolytics (Fig 1) and looks more similar to some of the antipsychotics (Fig 3), which could explain its frequent misclassification as an antipsychotic. Moreover, while the anxiolytic class had the fewest members (5), its members are also the most diverse as can be seen by the greater spread in the Tanimoto similarity coefficients (Fig 3).

### Planarian behavioral phenotyping

#### Activity and potency determined by benchmark concentration (BMC) modeling

The chemicals were screened acutely (< 3 hour exposure) in adult planarians and assessed for 13 endpoints spanning effects on lethality, body shape, stickiness, locomotion, and reaction to light and noxious heat (Fig 7, Tables 2 and 3). None of the tested compounds caused lethality at the tested concentrations, though a few compounds caused a substantial number of planarians to crawl out of their wells and dry out (“crawl-out behavior”). The BMC was calculated for each endpoint to determine when phenotypic responses exceeded empirically determined noise levels, defined by the benchmark response (BMR) (Fig 8, Tables 2 and 3, S10-11 Tables).

**Fig 7.**
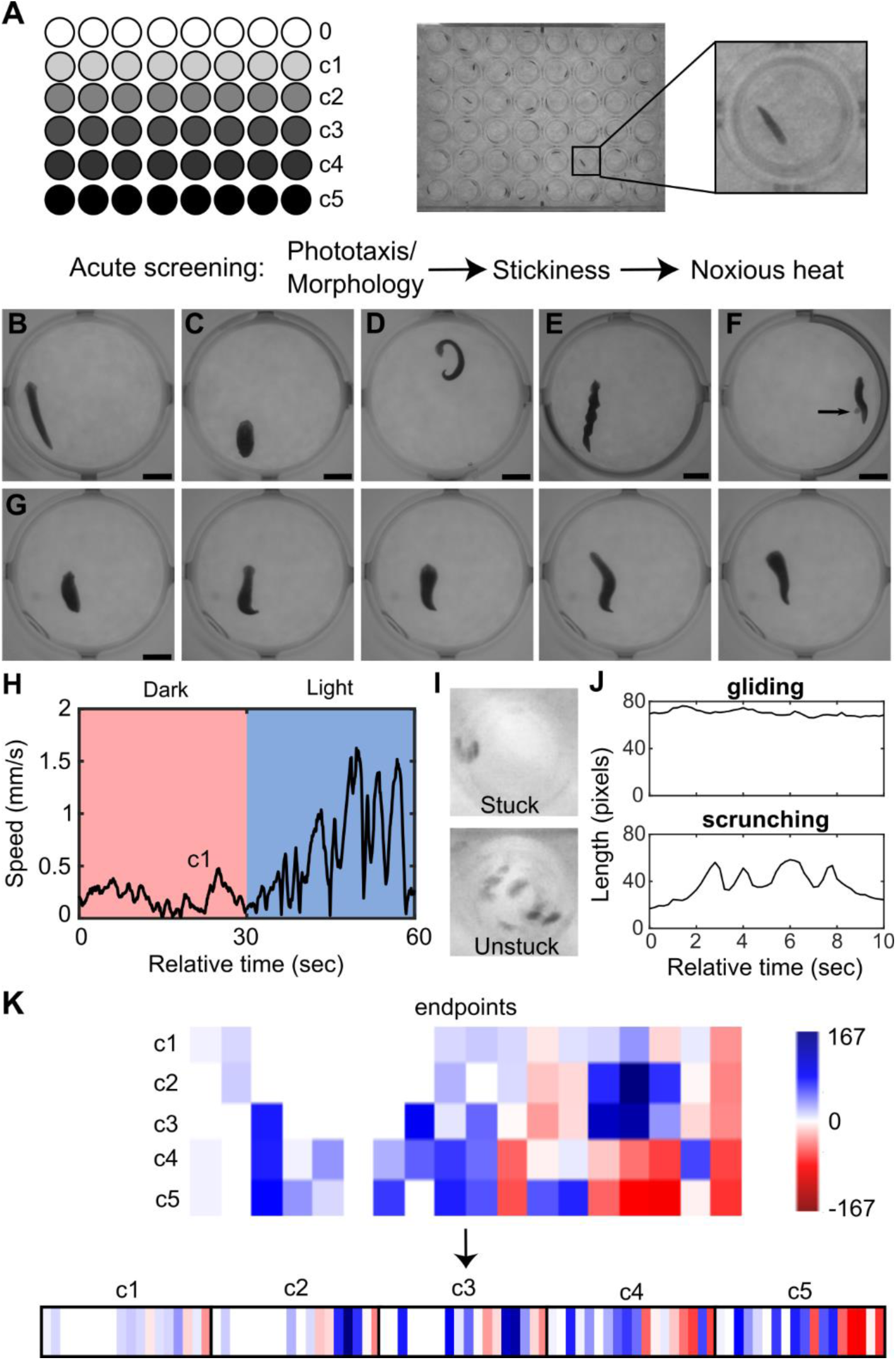
Overview of planarian screening and data analysis. A) Schematic of plate setup and representative image of a 48-well plate containing one planarian per well. Each plate tested a solvent control (c0) and 5 concentrations of a chemical and was tested acutely in phototaxis, stickiness, and noxious heat assays. B-G) Examples of various planarian body shapes: B) Normal planarian with smooth gliding. C) Contracted with shortened length and often associated with ruffling of the edges of the planarian. D) C-shape/curled and often on its side. E) Corkscrew showing multiple twists across the body axis. F) Pharynx extrusion. Arrow points at the unpigmented pharynx protruding from the underside of the planarian. G) Example image sequence of a planarian undergoing scrunching, showing oscillations of body length. This is one type of behavior, among others, which constitute the hyperkinesis category. Each frame is 1 sec apart. Examples shown are exposed to 0.5%(v/v) DMSO (B), 10 µM DRO (C and D), 10 µM BRO (E), 10 µM DUL (F), and 1 µM OLA (G). Scale bars: 2 mm. H) Representative plot of speed over time during the phototaxis assay. During a normal phototaxis response, planarians increase their speed during the blue light period. I) Minimum intensity projections of the shaking portion of the stickiness assay. A “stuck” planarian adheres to the bottom of the well and is not displaced, whereas an “unstuck” planarian dislodges and is displaced around the well. J) Length over time plots showing normal gliding or the oscillatory scrunching gait (49) that is induced during the noxious heat assay. K) Example schematic representation of the behavioral barcodes. First, a barcode is created for each chemical concentration, consisting of a numerical vector of the compiled normalized score for each endpoint (columns). One master barcode is made for each chemical by concatenating the barcodes for the individual concentrations. For details see Methods.

**Fig 8.**
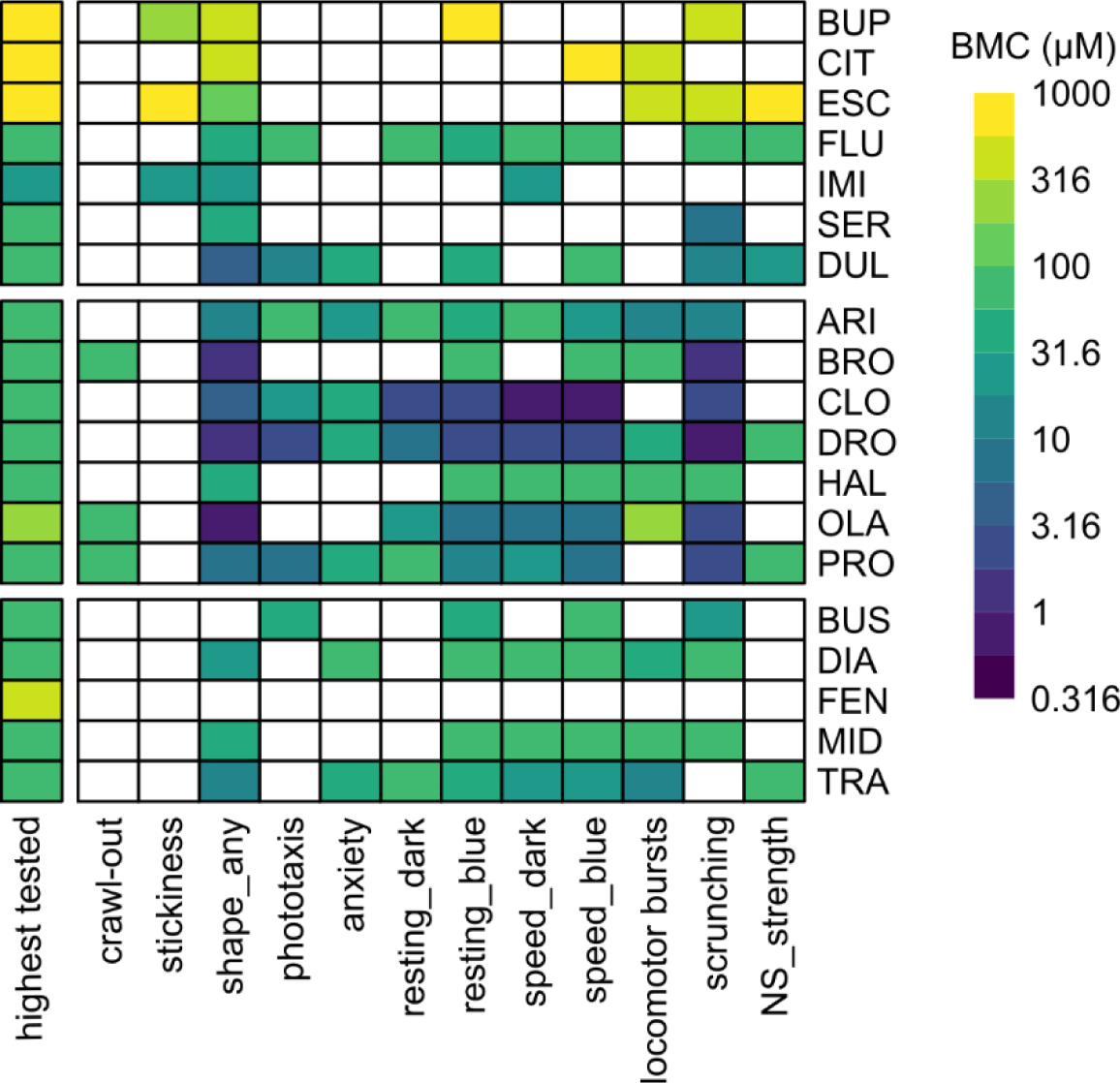
Activity of neuroactive drugs in adult planarians. Heatmap comparing the benchmark concentrations (BMCs) for all neuroactive compounds in adult planarians after acute exposure. The first column shows the highest tested concentration. As both “speed_blue” endpoints had similar BMC scores, only “speed_blue2” is shown as this was the more sensitive of the two timepoints. NS: noxious stimuli.

**Table 2.** Binary endpoints.

| Endpoint | Description | BMR<sup>a</sup> |
| --- | --- | --- |
| Crawl-out | % planarians that crawled out of the water | 25<sup>b</sup> |
| Stickiness | % stuck individuals | 25 |
| Shape (any) | % individuals with any abnormal body shape | 25<sup>b</sup> |
| Phototaxis | % planarians who did not phototax (50) | 35 |
| Scrunching | % non-scrunching planarians | 30 |

<sup>a</sup>BMR: benchmark response
<sup>b</sup>Variance was already minimized at all tested thresholds, thus the BMR was manually defined. See Methods.

**Table 3.** Continuous endpoints.

| Endpoint | Description | normalization | Direction<sup>a</sup> | BMR<sup>b</sup> |
| --- | --- | --- | --- | --- |
| Anxiety | Fraction of time spent in outer region of well | $(\text{Response}_{\text{chemical}} / \text{Response}_{\text{vehicle}}) * 100 - 100$ | - | 30 |
| Resting_dark | Fraction of time spent resting in 2<sup>nd</sup> dark cycle | $(\text{Response}_{\text{chemical}} - \text{Response}_{\text{vehicle}}) * 100$ | + | 60<sup>c</sup> |
| Resting_blue | Fraction of time spent resting in 2<sup>nd</sup> blue cycle | $(\text{Response}_{\text{chemical}} - \text{Response}_{\text{vehicle}}) * 100$ | + | 35<sup>c</sup> |
| Speed_dark | Mean speed (mm/s) in 2<sup>nd</sup> dark cycle | $(\text{Response}_{\text{chemical}} / \text{Response}_{\text{vehicle}}) * 100 - 100$ | - | 60<sup>c</sup> |
| Speed_blue1 | Mean speed (mm/s) in 1<sup>st</sup> 30 seconds of blue cycle | $(\text{Response}_{\text{chemical}} / \text{Response}_{\text{vehicle}}) * 100 - 100$ | - | 80<sup>c</sup> |
| Speed_blue2 | Mean speed (mm/s) in 2<sup>nd</sup> 30 seconds of blue cycle | $(\text{Response}_{\text{chemical}} / \text{Response}_{\text{vehicle}}) * 100 - 100$ | - | 70<sup>c</sup> |
| Locomotor bursts (total) | Sum of locomotor bursts in phototaxis assay | $\text{Response}_{\text{chemical}} - \text{Response}_{\text{vehicle}}$ | + | 20<sup>c</sup> |
| Noxious stimuli (strength) | Median displacement at end of noxious heat (47) | $(\text{Response}_{\text{chemical}} / \text{Response}_{\text{vehicle}}) * 100 - 100$ | - | 55<sup>c</sup> |

<sup>a</sup> The directionality (increasing (+) or decreasing (-) compared to the median control response) of the effect.
<sup>b</sup>BMR: benchmark response
<sup>c</sup>BMR was determined based on the 5<sup>th</sup> or 95<sup>th</sup> percentile of the histograms of the normalized vehicle control response (S2 Fig)

Except for FEN, all chemicals showed activity in at least one endpoint in at least one tested concentration. FEN did not show any observable effects up to 562 µM (the highest tested soluble concentration). Moreover, no lethality (0%, *n =* 32) or qualitative defects were observed in 562 µM FEN after a 12-day exposure (S5 Fig), suggesting it may not be bioavailable to planarians. ESC, the (*S*)-stereoisomer of CIT, was only active at ≥316 µM. However, OXA, its counterion control, also caused similar phenotypes at these concentrations (S12 Table). All other counterions for the salt forms of the drugs were found to be negative. To assess whether any of the observed effects were due to pH effects instead of compound-specific effects, the pH of the highest tested concentration of each chemical was measured (S13 Table). For reference, we measured the pH of Instant Ocean (IO) water as 6.72 ± 0.41 and that of 0.5% (v/v) DMSO as 6.63 ± 0.17 (mean ± SD, *n* =5), which is consistent with previous reports of acceptable pH ranges for planarian culture conditions (51). Acute behavioral effects were seen with acidic conditions at pH <4 (S13 Table). Both ESC and OXA had pH values <4 at the two highest tested concentrations (316 and 1000 µM), while at 100 µM, where no behavioral effects were observed, the pH was >4. Together, these data suggest that the effects seen at these concentrations may be due to low pH. Of note, 1000 µM CIT, which did cause behavioral effects, had a pH of 6.42, suggesting that the pH effects of ESC may have been driven by the OXA counterion. All other compounds were within acceptable pH ranges where no adverse effects were observed and thus the observed effects are presumed to be due to the pharmacological activity of the drugs.

Even within the same class, potency differences of several orders of magnitude were seen, which may reflect differences in uptake and metabolism or in affinity to the planarian neuronal targets. Lipophilicity can affect chemical bioavailability; therefore, we plotted BMC versus the calculated distribution coefficient, logD for each chemical. Because logD can vary depending on the pH, we plotted the range of logD values corresponding to the presumed pH range of the tested concentrations (ranging from the pH of the solvent as a proxy for the pH of the lowest concentration to the measured pH of the highest tested concentration (S13 Table)). We found no correlation between logD and BMC (S6 Fig), suggesting lipophilicity alone cannot explain the differences in potency.

Because of the observed potency differences, we evaluated the phenotypic profiles independent of concentration to identify any class-specific patterns (Fig 9). Defects in scrunching were seen in almost all the tested compounds, agreeing with our previous data showing that lack of heat-induced scrunching is a sensitive readout of disturbed neuronal function across a broad range of chemical types (44,46,48). Generally, the antipsychotics and anxiolytics showed a broad range of effects across multiple endpoints than the antidepressants (Fig 9A). For example, decreased motility in the dark and increased number of locomotor bursts were seen in >70% of the tested antipsychotics and anxiolytics, respectively, but only 29% of antidepressants. Effects in phototaxis and anxiety were more prominent in the antipsychotics than in the other classes. While “any abnormal body shape” was seen with almost all the tested drugs, the specific type of body shape differed across the drug classes (Fig 9B). For example, the antidepressants were largely characterized by the presence of hyperactive shapes/behaviors, without the inclusion of other body shape classes. In contrast, antipsychotics and anxiolytics had more mixed phenotypes with instances of contraction, C-shapes, and hyperactivity or C-shapes and hyperactivity, respectively.

**Fig 9.**
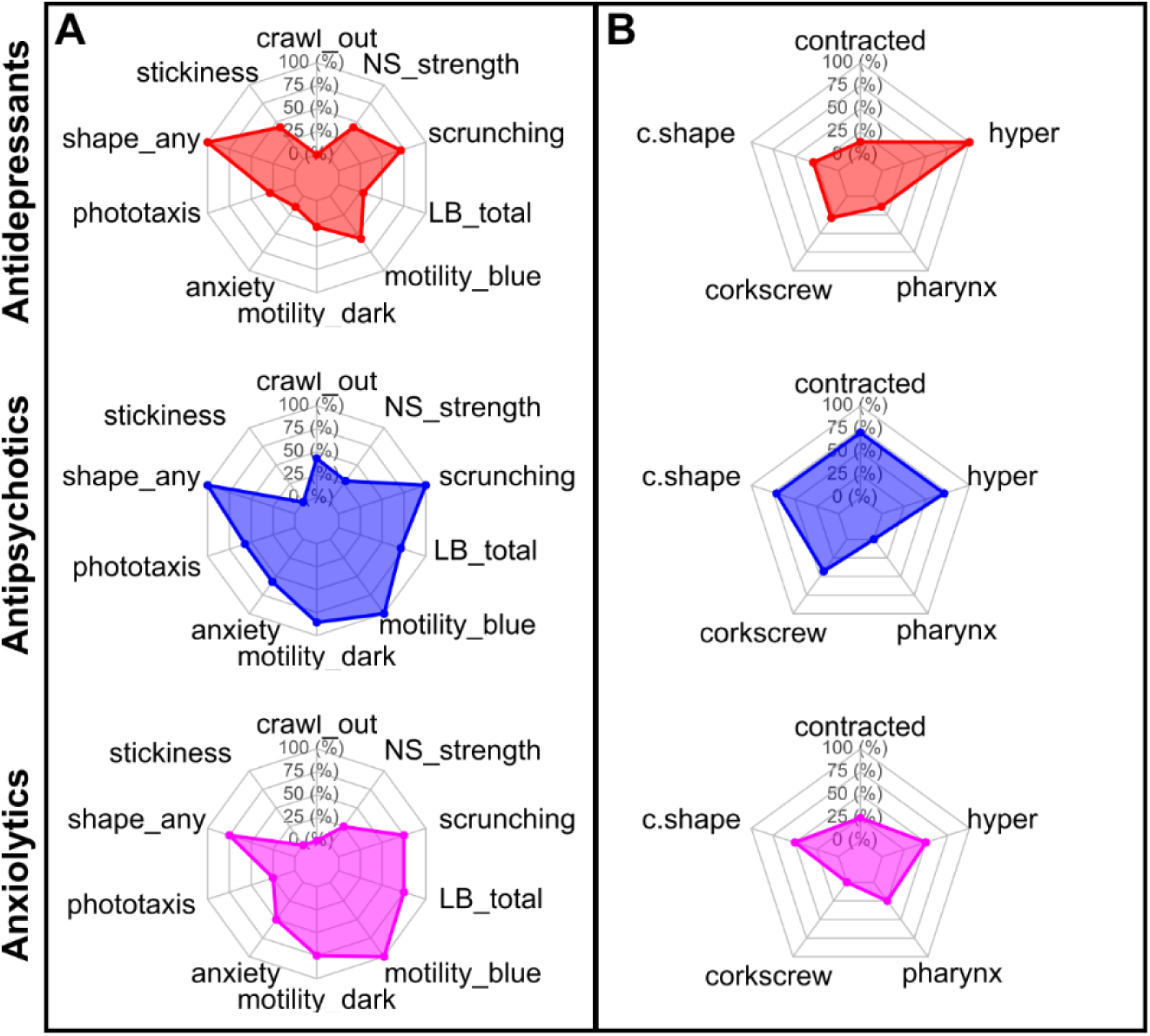
The drug classes showed different planarian phenotypic profiles. Radar plots showing the percentage of chemicals in each class that had a hit at any concentration in (A) any endpoint or (B) specific body shape classes. In (A) speed and resting endpoints in each light period were combined into a “motility” readout. LB: locomotor bursts; NS: noxious stimuli.

One possible confounding factor in this hit comparison, which aggregates effects at all concentrations, is that some of the highest tested concentrations may reach toxic levels and thus the observed phenotypes may be due to a mix of neuroactivity and toxicity. In fact, a common struggle in studies on neuroactive compounds is to distinguish neuroefficacy from toxicity, which both need to be considered during the drug discovery process. Importantly drug toxicity is the second most common reason (after lack of clinical efficacy) for promising drugs to fail in clinical trials (52); thus, it is important to identify possible toxicity early in the drug discovery pipeline. While we currently do not have sufficient understanding of the phenotypic readouts to be able to delineate neuroactive vs adverse effects, effects at low concentrations are likely to constitute efficacy and not toxicity. Because nominal concentrations do not necessarily reflect tissue concentrations due to compound-specific differences in uptake and metabolism, it is impossible to define a water concentration threshold that would separate efficacy from toxicity. Future work characterizing planarian drug metabolism will be helpful to gain a better understanding of how physicochemical properties affect chemical bioavailability. Therefore, in the absence of this knowledge, we included all nominal test concentrations for the planarian classifications.

#### Classification of neuroactive drug classes using planarian behavioral phenotyping

Ultimately, the strength of planarian behavioral screening is in the multidimensional information gained from looking across multiple endpoints testing various neurological functions and not just at each endpoint in isolation. To this end, each tested concentration of a compound was given a phenotypic barcode consisting of the compiled normalized score compared to in-plate solvent controls (see Methods, Figure 7K). For each chemical, one master barcode was created by concatenating the barcodes of all tested concentrations in relative order (S7 Figure).

To keep comparisons similar across different test concentrations, the relative order of the concentrations was considered rather than the absolute concentration. For the classification methods, the data were truncated to only the highest 5 concentrations such that negative/missing data were not driving the classifications. These planarian barcodes were then evaluated using the same computational methods as for the chemical features to determine how well the neuroactive drug classes could be separated (Figs 10-12).

**Fig 10.**
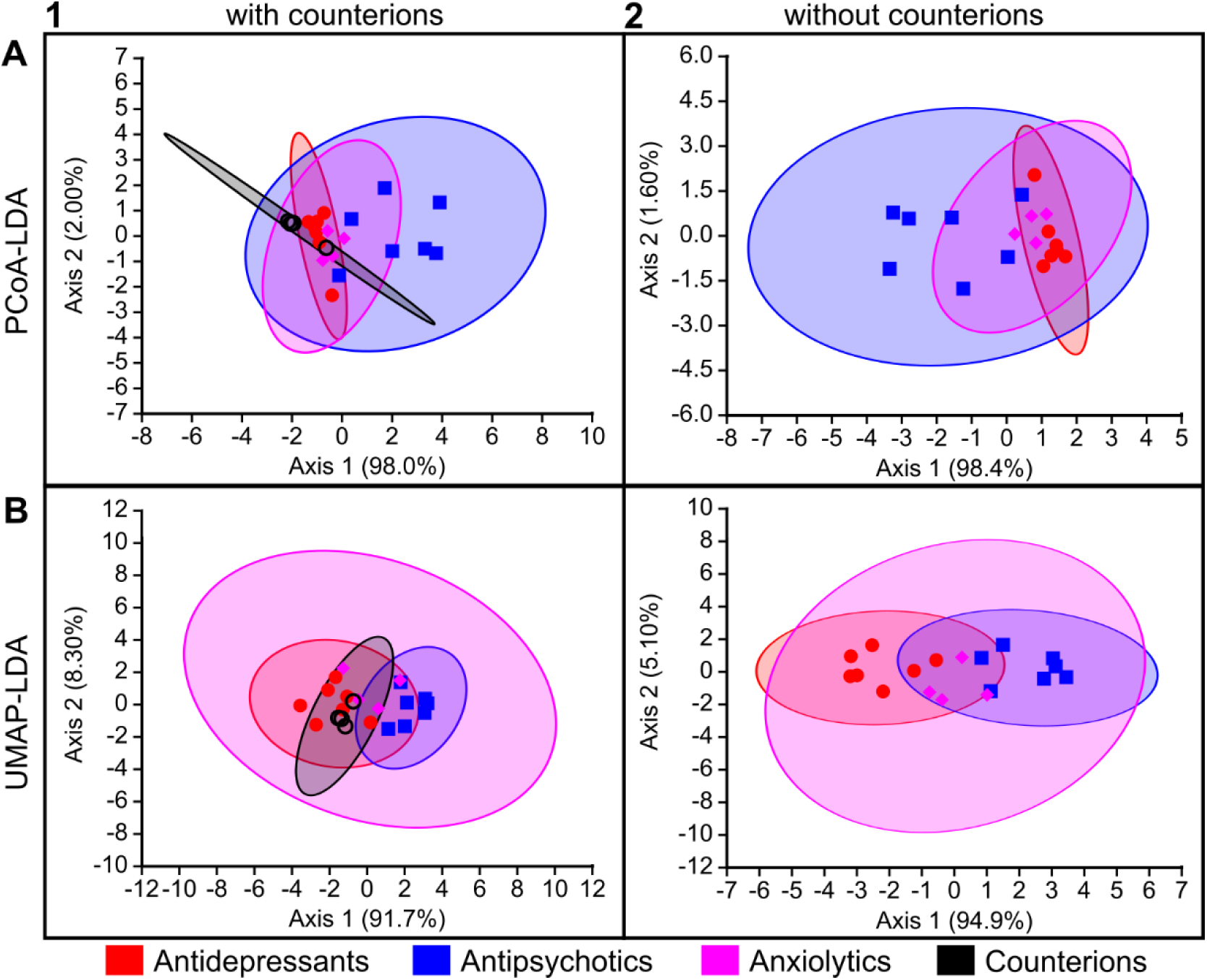
Classification of planarian behavioral phenotyping based on LDA-based approaches. Computational methods are arranged by rows: (A) PCoA-LDA; (B) UMAP-LDA. Counterion inclusion/exclusion is arranged by columns: (1) with counterions; (2) without counterions. Ellipses represent 95% confidence intervals. The axes show the percentage of the total eigenvalues.

**Fig 11.**
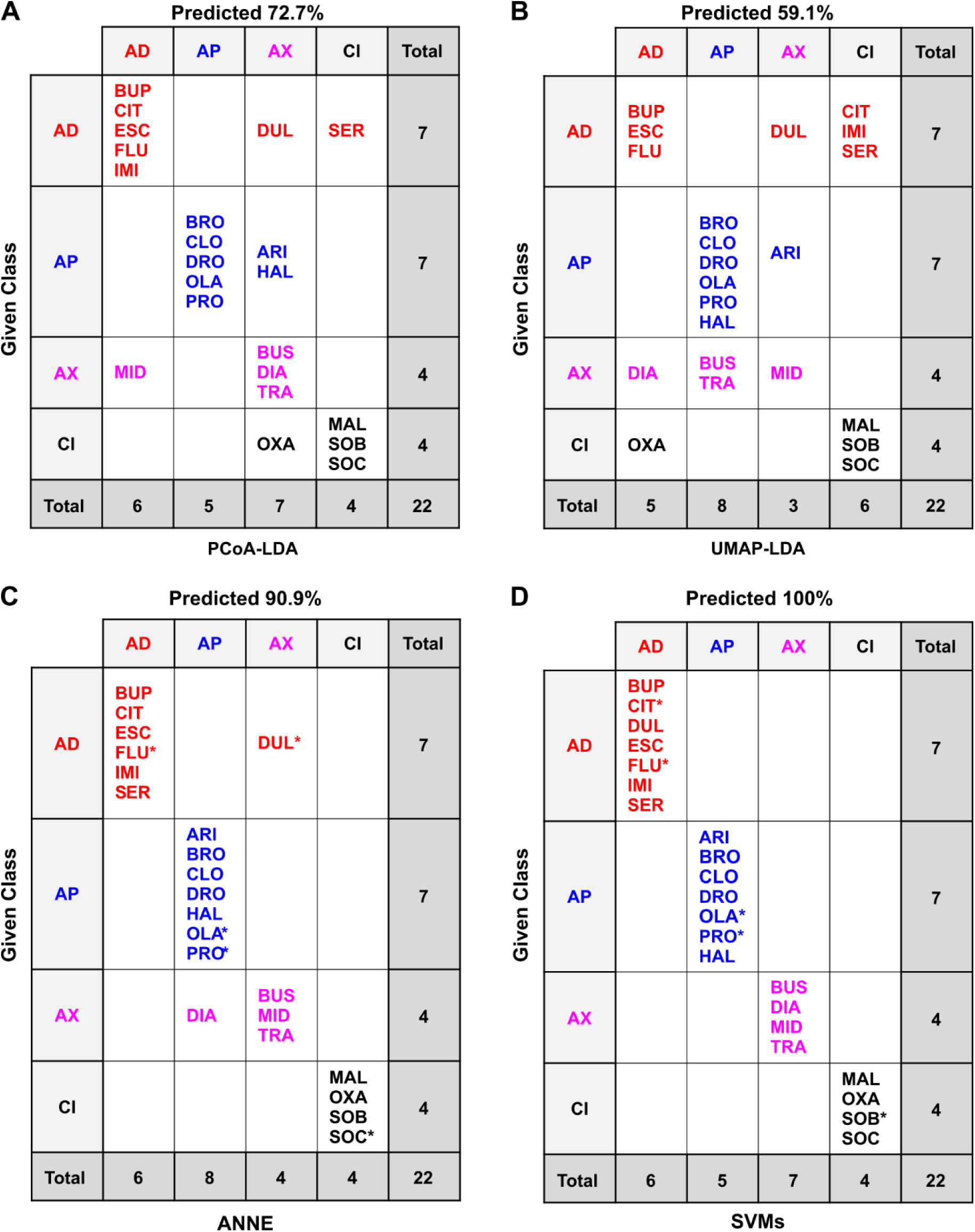
Confusion matrices of the classification methods based on planarian behavioral phenotyping of the drugs and counterions. Confusion matrices for the different classification methods: (A) PCoA-LDA; (B) UMAP-LDA; (C) ANNE; (D) SVMs. AD: antidepressant, AP: antipsychotic, AC: anxiolytic; CI: counterion. In A and B, predicted accuracy was calculated following an exhaustive jackknifing. In C and D, predicated accuracy refers to the overall accuracy. *indicates randomly chosen members of the test set. See the Methods for model selection. Test set accuracies were 80.0% for ANNE (S15 Table) and 100% for SVMs (S17 Table).

**Fig 12.**
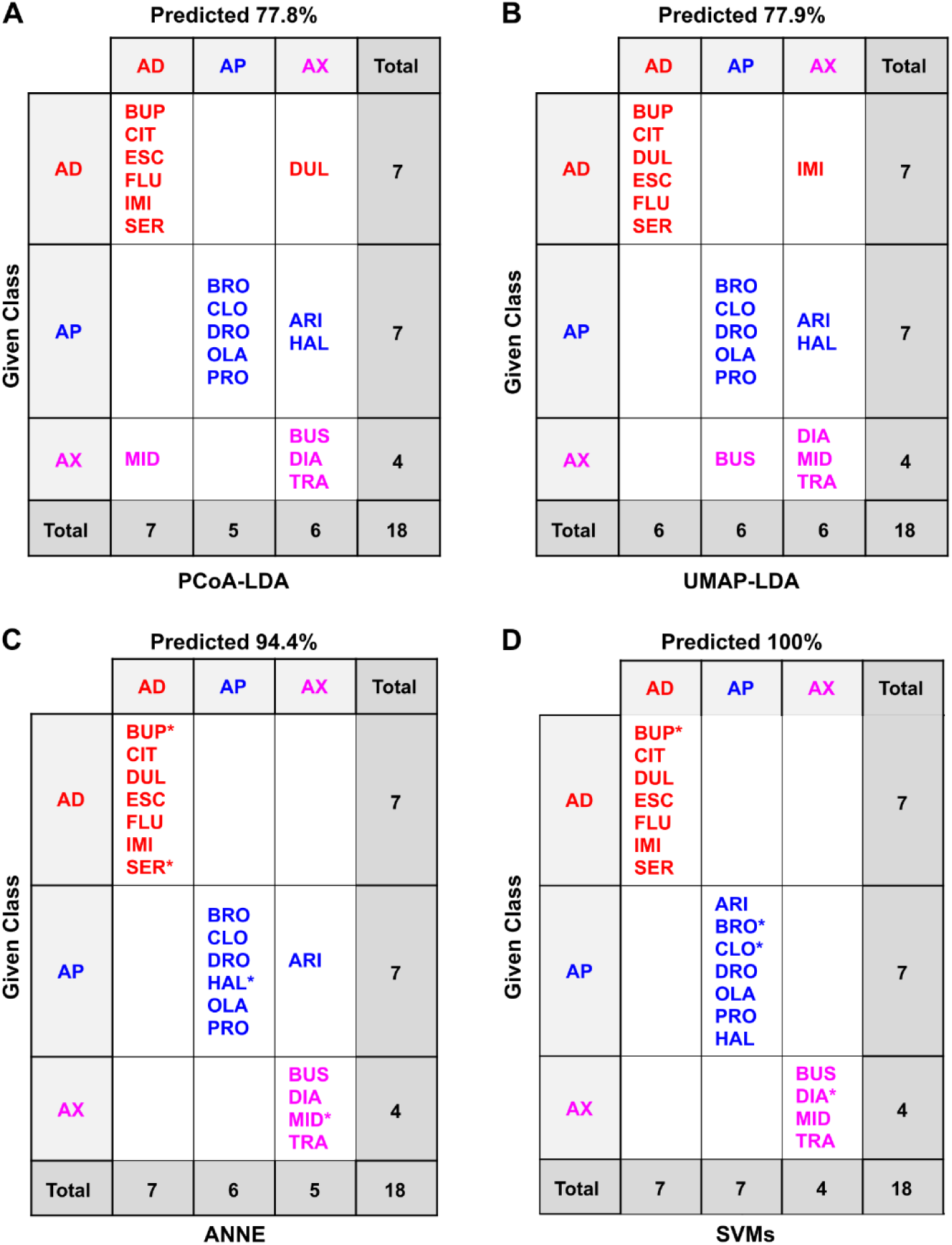
Confusion matrices of the classification methods based on planarian behavioral phenotyping of the drugs only. Confusion matrices for the different classification methods: (A) PCoA-LDA; (B) UMAP-LDA; (C) ANNE; (D) SVMs. AD: antidepressant, AP: antipsychotic, AC: anxiolytic; CI: counterion. In A and B, predicted accuracy was calculated following an exhaustive jackknifing. In C and D, predicated accuracy refers to the overall accuracy. *indicates randomly chosen members of the test set. See the Methods for model selection. Test set accuracies were 100% for ANNE (S16 Table) and 100% for SVMs (S17 Table).

As with the chemical descriptors, comparisons were made both with and without the counterions. ANNE and SVMs were each initially run across 10 different models (S15-18 Tables). The best performing model for each was chosen (see Methods) for comparison across methods. It is important to note that the algorithm used for training the ANNE models incorporated a stopping procedure to guard against overfitting, whereas the SVM algorithm did not provide this feature. However, while our process for selecting the best of ten models of each type included terms to reward accuracy, it also included terms to penalize complexity according to the number of neurons and descriptors for ANNE models or the number of descriptors for SVM models (see Methods for details).

All four classification methods were able to distinguish the neuroactive drug classes based on planarian phenotypic barcodes, with accuracies ranging from 59.1% to 100%. The machine learning methods, particularly SVMs, tended to perform better than PCoA-LDA or UMAP-LDA. Similar to the classification based on physicochemical data, UMAP-LDA performed the worst of the four methods. While this difference was clear when counterions were included (accuracy of 59% versus >72% in all other models (Fig 11)), UMAP-LDA and PCoA-LDA had practically the same adjusted accuracy when counterions were excluded (Fig 12). In contrast to the cheminformatic classifications, inclusion of the counterions decreased accuracy for all methods except SVMs, which had 100% accuracy in all cases. This decreased accuracy is likely because in both PCoA-LDA and UMAP-LDA the counterion OXA was “misclassified” as one of the drug classes due to the observed behavioral effects from the low pH.

Because FEN was inactive in the planarian screen, it was not included in these classifications. Inclusion of FEN (S8-10 Figs and S19-22 Tables) led to similar classification of the remaining drugs but showed overall lower accuracy due to the misclassification of FEN, either as a counterion or as an antidepressant. Notably, the lack of effects with FEN cannot be explained by its size or log D (S6 Fig).

Many of the compounds studied here have previously been tested in a behavioral screen in 7-day old zebrafish larvae (20). Similar to our findings for the classifications based on planarian data, the authors found that they could visually distinguish the three drug classes using multidimensional scaling, with considerable overlap of the clusters, particularly of anxiolytics and antidepressants (20). Given the lower complexity of planarians and their status as an invertebrate model (14), it was encouraging to find that we were able to perform effective behavioral classification schemes in a simple and cost-effective model.

### Planarian phenotyping adds information to cheminformatics

The goal of this pilot study was to determine the extent to which planarian behavioral phenotyping could provide additional information for neuroactive drug classification that could not be captured with cheminformatics alone. The value of using a combined cheminformatics and organismal HTS approach has been demonstrated for developmental toxicity and neurotoxicity studies using zebrafish larvae (53). Here, we show that planarian phenotypic data on neuroactive chemicals augments the cheminformatics data.

The classification of the chemicals based on the planarian behavioral phenotyping did not use any information about the chemicals. Thus, the fact that classification accuracy reached > 70% for all models (except for UMAP-LDA with counterions included) and >90% for some models for the planarian data suggests that behavioral barcodes contain meaningful information that reflects the underlying biology. Moreover, while PCoA-LDA and UMAP-LDA classification methods had higher accuracy when using chemical descriptors rather than planarian behavioral barcodes, the machine learning models had similarly high accuracies (>90%) for both data streams. Thus, the choice of classification method can be equally as impactful as the choice of input data. Taken together, our data imply that planarian behavioral screening can identify and differentiate similarly acting compounds without knowledge of the underlying biology or physicochemical properties, making it well suited for unbiased, discovery-based screening of neuropsychiatric drug candidates.

Some of the most misclassified compounds using behavioral responses across the different methods included DUL and ARI (Fig 11 and 12). Both compounds have known dual functions. DUL affects both serotonin (5-hyroxytryptamine; 5HT) and noradrenaline reuptake (54), and while it is mainly classified as an antidepressant, it has been shown to have anxiolytic effects upon chronic exposure in mice (54,55). Across all classification methods except for SVMs, DUL was often classified as an anxiolytic. Similarly, ARI was originally classified as an atypical antipsychotic but is also used clinically to augment antidepressant efficacy (56,57). When using planarian phenotyping, ARI was often classified as either an antidepressant or anxiolytic. Thus, planarian phenotyping was able to detect known “off-label” effects of these drugs that could not be detected by cheminformatics alone. Moreover, planarian behavioral barcoding correctly identified BUP as an antidepressant in all models – in contrast to our findings with the physicochemical data. BUP is an aminoketone and its mode of action as a norepinephrine and dopamine reuptake inhibitor is distinct from that of the other antidepressants which are all selective serotonin reuptake inhibitors (58). This knowledge of the *in vivo* effects can only be gained from an organismal systems-level approach, where the effects of these pathway interactions can be observed.

### Study limitations and considerations for follow-up studies

This pilot study was small in scope (19 compounds) and faced limitations that need to be considered when designing follow-up studies with larger compound libraries to support or refute the findings reported here. First, we tested only 5 anxiolytics because most are controlled substances, and thus more expensive and difficult to obtain and use than other compounds. Furthermore, given that FEN was inactive in planarians up to the solubility limit, this reduced the anxiolytic category to only 4 active compounds for the planarian classifications compared to 7 compounds in each of the other two categories. While the numbers of compounds were still comparable here, class imbalance is something to consider when screening larger libraries (59,60).

For planarian behavioral phenotyping, we created an aggregated barcode that included effects across all tested concentrations, thus potentially including toxic effects at higher concentrations for some chemicals. By looking at the whole concentration profile for a chemical, we anticipated finding phenotypic signatures for compounds that act similarly, but it is possible that the effects were not aligned by relative concentration or that toxicity of different chemicals can manifest in different ways and confound the classifications. Some zebrafish studies have overcome this issue by selecting single representative concentrations of a chemical to use as a comparator (20). However, it is not always straightforward to determine which concentration is truly representative and reflective of substantial neuroefficacy without the confounders of toxicity.

We evaluated four computational models and found them to have comparable performance on our data, which made it difficult to decide on a “best” classification method, likely due to the small scope of this study. The machine learning models reached >90% for all cases, thus generally performed better than the LDA methods, but they differed in misclassifications and information content. The LDA methods provide insight into the similarity/distances between chemicals which allows for visualization of the classification (Figs 4 and 10) and are not provided by the ANNE and SVMs, which are more enigmatic in how the models create the classifications.

The next steps to further test these classification methods would be to screen larger libraries of neuroactive compounds that are commercially available, as for example in (20,22). Having a larger library would allow for a truly separate and larger test set of chemicals to evaluate model performance instead of the jackknifing approach conducted here. For these initial scaling-up efforts, it will be important to select chemical libraries that contain several members of the same category while also spanning different classes and to include chemicals with both known and unknown modes of action.

One consideration for testing larger libraries of 1000+ chemicals is feasibility. While small organism HTS is substantially faster than vertebrate testing it is still an experimental method and thus will be rate-limiting compared to a purely computational approach. To screen larger libraries, one could select one or two concentrations based on the tests conducted in this pilot study for an initial screen. Further screening for concentration response information could then be performed on promising candidates from the initial screen. This approach has been successfully used in zebrafish larvae to conduct such large library screens (20,22). Moreover, the increased incorporation of robotic handling systems will expand the throughput capabilities of small organismal screens.

Upon generating larger datasets, the next step would be to integrate the two data streams used here (cheminformatics and planarian phenotyping) to generate a single classification. Other available data streams, such as those generated from *in vitro* HTS (61,62) or computational modeling of drug-target interactions (63,64), could also be integrated, similar to the Integrated Approaches to Testing and Assessment (IATA) framework used in toxicology to combine multiple data streams to assess chemical toxicity (65,66).

A few recent studies have implemented such a synthesis based on different types of information, e.g., cheminformatics, and/or HTS data, including *in vitro* cell painting and gene expression profiles, e.g. (67,68). How the data streams are combined and weighed is a non-trivial decision that will itself require validation with known chemical libraries. Thus, there are multiple steps to take and decisions to make that need to be carefully considered and scrutinized to build upon the results from this pilot study. The important contribution of this work is that it has shown the feasibility and promise of invertebrate behavioral HTS as a viable experimental alternative to vertebrate testing for first-tier screening of novel drug candidates.

## Materials and methods

### Chemicals and chemical structures

Neuroactive compounds that have been functionally classified as antidepressants, antipsychotics or anxiolytics were studied. Functional classifications were assigned based on the primary classification indicated on the manufacturer’s website and/or previous literature (20). Table 1 lists the chemicals studied including the chemical abstracts service (CAS) numbers and information pertaining to the use of the compounds in the experimental studies. Some chemicals were provided in salt form; thus, we also tested the respective counterions alone (Table 1).

For computational models based on two-dimensional (2D) chemical structures of drugs and counterions, 2D coordinates were downloaded as structure-data format (SDF) files from PubChem (https://pubchem.ncbi.nlm.nih.gov/) and concatenated into a single 2D SDF file using OpenBabel 3.1.1 (https://github.com/openbabel/openbabel) (69) for Linux (Linux Mint 21.3 Xfce, https://linuxmint.com/). For computational models based on three-dimensional (3D) chemical structures of drugs and counterions, 3D coordinates were downloaded as SDF files from PubChem, energy-minimized in the AMBER14 force field at pH 7.40 (Krieger et al., 2006, PMID: 16644253; Maier et al., 2015, PMID: 26574453) using YASARA-Structure 23.12.24 (Ozvolkik et al., 2023, PMID: 37782001) for Linux and exported as a single concatenated 3D SDF file. Version 2000 was used for both 2D and 3D SDF files. The ionization state of each compound at pH 7.40 was confirmed using the pKa module of Marvin Sketch 23.1.0 for Linux (https://www.chemaxon.com). Structures, names, 3-letter codes, and primary functional pharmacological classes of the drugs and counterions used in the computational studies are shown in Figure 1.

### Chemical similarity

Chemical similarity of the 3D SDF structures of drugs and counterions relative to ARI as a reference structure was assessed using vROCS 3.6.1.3 (OpenEye, Cadence Molecular Sciences, Santa Fe, NM, https://www.eyesopen.com; (63)) for Linux. Similarity was expressed quantitatively in terms of the Tanimoto coefficient (Tc), which is the most widely used similarity metric in cheminformatics (70,71). The program calculates three Tc values: Tc Shape, based on a 3D alignment of two molecules that maximizes the volume of overlap; Tc Color, derived from a 3D alignment of two molecules with respect to six molecular features (H-bond donor, H-bond acceptor, anion, cation, hydrophobe, and ring); and Tc Combined, the sum of Tc Shape and Tc Color. The numerical values of Tc Shape or Tc Color range from 0 (no similarity) to 1 (complete similarity); therefore, Tc Combined ranges from 0 (no similarity) to 2 (complete similarity). Note that two identical molecules would have Tc Shape and Tc Color coefficients of 1. However, if two structures have Tc Shape and Tc Color coefficients of 1, this does not necessarily mean that the molecules are identical because two non-identical compounds could have distinctive features that were not accounted for in the similarity algorithms (72).

### Chemical descriptors

Chemical descriptors (comprising computed chemical properties and fingerprints) were calculated for 2D and 3D structures of drugs and counterions using absorption, distribution, metabolism, excretion, and toxicity (ADMET) Predictor^®^ 11.0.3 (Simulations Plus, Lancaster, CA) for Windows (10 Pro 22H2). The fingerprint type was extended connectivity fingerprint diameter 6 (ECFP6). Within ADMET Predictor^®^, fingerprints were converted to descriptor format and combined with the other computed chemical properties descriptors for use in the computational models. For each of the 2D structures (18 drugs and 5 counterions) there were 908 descriptors (514 fingerprints and 394 directly calculated properties), and for each of the 3D structures (21 drugs and 5 counterions) there were 948 descriptors (516 fingerprints and 432 directly calculated properties). Importantly, when considering 2D structures CIT and ESC are treated as one structure which we label as CIT, leading to only 18 drugs. Because 3 of the 18 drugs were administered as racemic mixtures in the behavioral assays, the separate structures of the (*R*) and (*S*) stereoisomers of these compounds were included in the 3D computational models, thereby giving rise to 21 3D drug structures (and 5 counterions). The 2D and 3D descriptors were exported from ADMET Predictor^®^ as two separate data files in CSV format for use in classification models. To keep the data size comparable to the planarian data, which considers each of the 19 tested compounds as one entity, we focused on the 2D structural features in the main text.

### Classification models

#### Linear discriminant analysis (LDA) preceded by principal coordinate analysis (PCoA)

LDA is a supervised linear ordination and classification technique that maximizes the discrimination between classes of labeled data (73). LDA is a preferred classification procedure when there are more than two classes, in which cases the method is also known as canonical variate analysis (CVA) or canonical discriminant analysis (CDA). When the number of predictor variables is greater than the number of samples, LDA is preceded by a dimension reduction technique such as principal component analysis (PCA) or PCoA (74).

We chose PCoA as the dimension reduction step because PCoA can use any desired distance or dissimilarity measure, although Gower distance tends to be preferred (75), whereas PCA is restricted to Euclidean distance. Moreover, PCoA has been combined with LDA to create a successful sequential method for ordination and classification of standardized data called canonical analysis of principal coordinates (CAP) or generalized discriminant analysis (GDA) that preserves the initially chosen distance or dissimilarity measure in the resulting ordination (76).

The procedures for PCoA-LDA were carried out as follows: Either the chemical descriptor files or the barcode files of planarian behavioral responses were imported into the data analysis program, PAST 4.16c for Windows (https://www.nhm.uio.no/english/research/resources/past/; (77,78)). Classifications based on 2D or 3D chemical descriptors were carried out with or without the counterions. Classifications based on planarian responses were conducted on data with or without FEN and with or without counterions. Columns that were deemed uninformative by the PAST software (non-numeric, all-zero, all-missing, singleton, and constant) were removed, and the remaining data were standardized so that each column had mean = 0 and SD =1. Following dimension reduction in PAST using PCoA with Gower distances (79–81), LDA was carried out in PAST using the number of PCoA coordinates that yielded the highest classification accuracy (expressed as the percentage of correctly classified chemicals) after applying an exhaustive leave-one-out procedure (jackknifing) (82,83). Clusters identified by LDA were displayed as 2D scatterplots with the percentage of the total eigenvalues shown on each of the two axes and the clusters demarcated as 95% confidence ellipses. When the number of PCoA coordinates used for LDA was greater than two, the sum of the percentages of the total eigenvalues on the two axes was less than 100%.

#### LDA preceded by uniform manifold approximation and projection (UMAP)

UMAP was carried out on standardized data files based on chemical descriptors or planarian behavioral responses in PAST 4.16c using Gower distances as described above for PCoA. The UMAP algorithm employed by PAST was derived from the original method of (84). UMAP is a nonlinear dimension reduction and ordination method that has capabilities for preserving both the local and global structures of the original data, as an alternative dimension reduction method prior to LDA. UMAP provides a complementary alternative to PCoA, which is a linear dimension reduction and ordination technique that emphasizes preservation of the global structure of the original data (85).

The numbers of embedded and UMAP neighbors, as well as the minimum distance between samples, were systematically varied to achieve the maximum separation of clusters after 100 UMAP iterations. Final UMAP coordinates were then used as input for running LDA in PAST as described above for PCoA-LDA. The resulting LDA output included 2D scatterplots with 95% confidence ellipses and percent classification accuracies (unadjusted and jackknifed).

#### Artificial neural network ensemble (ANNE)

An artificial neural network (ANN) is a mathematical construct that mimics a simple array of biological neurons consisting of an input layer, a hidden layer, and an output layer (86). Inputs with various weights are transmitted to neurons in the hidden layer, where the summed weighted inputs are compared with a threshold value generated by a sigmoid activation function. When the threshold value is exceeded, a signal is conveyed to the output layer. ANNs have the capacity to accept nonlinear and large numbers of inputs from which they can adapt and learn, thereby enabling them to make predictions in the form of continuous outputs (regression models) or discrete assignments to two or more categories (classification models) (87). Multiple ANNs can be combined into an ensemble (ANNE) so that their outputs are channeled to a voting or averaging algorithm to create an ensembled output, resulting in enhanced performance of ANN-based regression and correlation models (88).

ANNE classification was performed using the Modeler™ 11.0 module of ADMET Predictor^®^ 11.0.3, with CLASS as the dependent variable. The classes consisted of the four primary functional categories (antidepressants, antipsychotics, anxiolytics, and counterions). Ten final models were created for each of the following four sets of chemical descriptors: 2D descriptors with or without counterions; and 3D descriptors with or without counterions.

Likewise, ten final models were created for each of the following four sets of planarian behavioral data: including FEN with or without counterions; and excluding FEN with or without counterions.

Before generating the models, the program screened all the descriptors to remove those with the following characteristics: identical or with coefficients of variation ≤ 1%; under-represented (non-zero for 1-3 data points); or highly correlated (Pearson *r* ≥ 0.98). For chemical descriptors, the culling process reduced the number of candidate 2D descriptors from 908 to 162 and the number of candidate 3D descriptors from 948 to 192. During the running of the ANNE and SVMs models, the algorithms selected the smallest number of descriptors that optimized performance; for some ANNE models, the final number of descriptors was as low as one, and for some SVMs models, the final number of descriptors was as low as two. S1 Table is a compilation of all of the chemical descriptors that were used in constructing the ANNE and SVM models listed in S2-S9 Tables. For planarian behavioral descriptors, the culling process reduced the number of candidate descriptors from 87 to 80 (FEN with counterions), 87 to 81 (FEN without counterions), 87 to 80 (without FEN with counterions), and 87 to 81 (without FEN and without counterions).

The training:test ratios of chemicals for chemical descriptor files were as follows: 2D with counterions, 18:5; 2D without counterions, 14:4; 3D with counterions, 21:5; and 3D without counterions, 17:4. These ratios for planarian behavioral files were as follows: FEN with counterions, 18:5; FEN without counterions, 15:4; without FEN with counterions, 17:5; without FEN without counterions, 14:4. Test set selection was done via randomized stratified sampling by CLASS. During each run, the members of the ensemble automatically were selected by initially assigning random weights to each submodel and partitioning the training sets into training and verification sets in an approximate 2:1 ratio. After training, scores were assigned to each model according to the summed verification set performance and the absolute difference between verification and training set performance using the Youden index as the criterion. To avoid overtraining of ANN models, training was automatically halted when the verification set score failed to improve or increased beyond a preset number of iterations.

Generation of each of the 10 final models was initiated using a unique random seed number. Final descriptor selection for each model was carried out by the software using a genetic algorithm to explore the effectiveness of different descriptor combinations. Models were then run using default settings for the number of generations, number of neurons, and number of descriptors; these settings were automatically adjusted to be appropriate for the number of chemicals in the training and test sets.

For each run, the following performance metrics generalized to *k* classes and expressed as percentages were calculated: Youden index (*J*) (89), Matthews correlation coefficient (MCC or *R*_k_) (90), and Accuracy (Acc) (overall percent correctly classified) (Farhadpour et al., 2024). These metrics were calculated by the Modeler™ 11 software for the training set, test set, and all chemicals. The data summaries include the following information for each model: mean and SE (*n* = 10 models) for each of the 9 performance indicators; number of neurons and descriptors; and the numbers and identities of misclassified chemicals. The rank for each ANNE and SVMs model, respectively, was determined by applying the RANK.AVG function in Microsoft Excel 365 to SUM(training metrics + test metrics + (100×*N*_min_/*N*) + (100×*D*_min)_ /*D*)), where *N*_min_ = minimum number of neurons, *N* = number of neurons, *D*_min_ = minimum number of descriptors, and *D* = number of descriptors. Thus, the ranking rewarded models that produced high performance with the smallest numbers of neurons and/or the smallest numbers of descriptors, in keeping with the parsimony principle (91,92)

#### Support vector machines (SVMs)

This is a machine-learning technique that seeks to define hyperplanes in *n*-dimensional space that maximize the separation of data points into classes (93). The name of the method stems from the fact that the points closest to a given hyperplane are called support vectors (94). Although SVMs can be used for regression models and outlier detection, they are especially well suited for classification (95). SVMs are intrinsically a binary classifier, but they have been successfully adapted for classifications involving multiple classes.

SVMs classification was performed using the Modeler™ 11.0 module of ADMET Predictor^®^ 11.0.3, with CLASS as the dependent variable, as described above for ANNE classification. Likewise, the same evaluation statistics and data summaries employed by ANNE were used by SVMs.

### Planarian Care and Culture

*D. japonica* planarians from an established lab culture were used for all experiments and cultivated according to standard protocols (44,47). The planarians were kept in 0.5 g/L Instant Ocean (IO) Salts (Spectrum Brands, Blacksburg, VA, USA) in BPA-free polypropylene plastic containers (approximately 25 cm L x 14 cm L x 8 cm H), with the lid on loosely and stored at 20°C in a Panasonic refrigerated incubator in the dark when not used for experiments. The planarians were fed 1-2x per week with 100% grass-fed beef liver (purchased from a local farm) or USDA-certified organic chicken liver (Bell and Evans) and cleaned on the feeding day and 2 days later. When not fed, the containers were cleaned weekly. Similarly sized intact planarians that were fasted for 5-7 days were arbitrarily selected to be used in experiments, as in (48).

### Chemical exposure

Stock solutions were prepared at 200X of the highest test concentration in either DMSO (Sigma-Aldrich, purity = 99.9%) or Milli-Q water, depending on solubility (Table 1). The 200X stocks were diluted to 10X stocks in IO water just prior to exposure. Planarians were exposed to the chemicals in 48-well tissue culture-treated polystyrene plates (Genesee Scientific, San Diego, CA, USA), with each well containing 1 planarian in 200 µL of chemical solution. When used as a solvent, DMSO was used at a final concentration of 0.5% (v/v), which does not cause morphological or behavioral effects in *D. japonica* (43).

For most chemicals, serial half-log dilutions were used to prepare the range of concentrations tested, which were initially guided by previous results in developing zebrafish (20) or our previous studies with *D. japonica* (96). For FEN and MID, serial quarter-log dilutions were used because preliminary tests had already narrowed an appropriate concentration range to span no effects to strong effects (or maximum solubility). All chemicals were screened over at least 5 concentrations, with each row of the 48-well plate consisting of one concentration (*n* = 8). For some compounds, additional lower concentrations were screened if the original lowest concentration showed effects in the preliminary data analysis. Moreover, some compounds were screened at additional higher concentrations if no statistically significant effects were observed in the original range and the compound was still soluble at higher concentrations. The counterions were screened at the concentration equivalent to what would be found in the highest concentration of the respective salt form of a tested drug. The pH of the highest test concentration for each chemical was first measured with pH strips (VWR, Radnor, PA). As we found the measurements with the pH strips were not precise or reliable, we remeasured the pH using an Apera PH60-MS pH Tester kit for small volumes (Apera Instruments, Columbus, Ohio), taking measurements after allowing the probe to equilibrate for 20 min.

Plates were sealed with thermal film (Excel Scientific, Victorville, CA, USA) immediately after addition of chemicals. Experiments were run on intact planarians within 15 minutes – 3 hours of chemical introduction. This time window was chosen for throughput, to allow for multiple plates to be set up and tested in one experiment. Which plates were screened first was arbitrarily decided, to avoid any systematic bias in the treatment. Technical triplicates were run for all chemicals (total *n* = 24 per concentration), employing a rotating orientation of the chemical concentrations in the plate rows to account for edge effects when screening (44).

One screening plate consisted of negative and positive assay controls, each run at a single concentration, which were used to evaluate proper performance of the assays. Negative controls consisted of D-sorbitol and L-ascorbic acid (both from Sigma-Aldrich) at 100 µM (44). Ethanol (1% v/v, Greenfield Global, Toronto, Canada), DMSO (3% v/v), and sodium dodecyl sulfate (SDS, 1 mg/L, Life Technologies, Carlsbad, CA) were used as positive controls at concentrations that induce behavioral phenotypes in the absence of lethality in *D. japonica* (43). The positive controls were prepared fresh at 10x in IO water on the day of the experiment. All assay controls performed as expected (S23 Table).

### Planarian screening methodology

Intact planarians were assayed for acute effects on lethality/crawl-out behavior, morphology, stickiness, and various behaviors (locomotion, phototaxis, and noxious heat sensing) using the custom screening platform and analysis methodology described in (44,47,48,50). Image analysis was performed using custom scripts in MATLAB or Python. Because the liquid volume only takes up 13% of the total well volume, planarians can crawl out of the liquid and dry out. Planarians which crawled out of the water and died (crawl-out, also previously referred to as “suicide” (44)) were excluded from the analysis of all other endpoints. Some planarians crawled out of the water during screening; these worms were included in the analysis for assays in which they were still alive.

The endpoints used are described in Tables 2 and 3 and consisted of two major classes: unstimulated and stimulated behaviors. Unstimulated behaviors included lethality/crawl-out, presence of abnormal body shapes/behaviors, and general locomotion (speed and resting in the dark). Specific distinctive body shape categories (contraction, C-shape, corkscrew, pharynx extrusion, and hyperkinesis) were manually scored blind by a researcher (Fig 7B-G).

Contraction, C-shape, and corkscrew were scored as described in (96). Hyperkinesis consisted of an array of different types of generally hyperkinetic movements that could not be captured by the other categories, such as muscle waves/scrunching (96), hyperextension, head flailing and convulsive behavior, consisting of uncoordinated muscle twitching (S1 Movie). A single planarian could be scored as exhibiting up to three different shape categories. For benchmark concentration (BMC) modeling, described below, the presence of any abnormal body shape was used to determine activity, whereas for phenotypic profiling, both incidence rate of any body shape and the incidence rates in the specific shape classes were used. Stimulated behaviors were measured in response to an environmental stimulus (shaking, light, or noxious heat, Fig 7).

### Benchmark concentration modeling

Benchmark concentrations (BMCs) were calculated for every chemical and endpoint to quantify potency using the Rcurvep R package (97) similar to the procedure described in (48). Briefly, the planarian responses for each endpoint were transformed as needed to allow for determination of directional, concentration-dependent responses, as described in (48). For the binary endpoints (crawl-out, body shape, stickiness, phototaxis, scrunching), the incidence rates (number of planarians affected and total number of planarians) from the combined data from all replicates (*n* = 24) was used. For all binary endpoints except crawl-out, the experimental incidence rates for each concentration were normalized by the incidence number of the respective in-plate vehicle controls. Any negative incidence numbers after normalization were set to 0. For continuous endpoints, the raw response of each individual planarian was normalized either by dividing by or subtracting the median of vehicle control values for that plate (Table 3). Except for locomotor bursts (total), the normalized outcome measures were multiplied by 100 to represent the percent change from the control populations and to provide an appropriate range to perform the BMC analysis. The normalized data were used as input for the Rcurvep packaged to calculate the benchmark response (BMR) as described in (48). For some endpoints, the R package could not converge to produce an accurate BMR due to high variance and/or instance of non-monotonic concentration responses. In these cases, the BMR was set to the 5^th^ or 95^th^ percentile of the normalized response of the aggregated vehicle controls (S4 Fig), to set the threshold above which effects would be seen. The determined BMR was then used to calculate the BMC, the concentration that exceeds that BMR in the modeled concentration response curve (97), for each endpoint, using *n* = 1000 bootstrapped curves. For all endpoints, we report the resulting median BMCs from this bootstrapped analysis. The lower and upper limits (5th and 95th percentiles, respectively) of the BMC for each endpoint are listed in Supplemental File 1. Some endpoints can be affected in both directions (e.g., increases or decreases in speed). For these endpoints, BMRs and BMCs were calculated for each direction, but we found that hits were only determined in one direction and thus only report that direction (Table 3).

### Phenotypic profiling

To quantitively describe the multidimensional phenotype observed for each test concentration, a “phenotypic barcode” was created for each chemical concentration consisting of either the incidence rate for all binary endpoints, multiplied by 100, or the median normalized response for all continuous endpoints, as were input into the BMC analysis. Next, one master barcode was created for each chemical by concatenating the barcodes of all tested concentrations in relative order. To keep comparisons similar across different test concentrations, the relative order of the concentrations was considered rather than the absolute concentration. Because different numbers of concentrations were tested across the different chemicals, we “right-aligned” the barcodes such that the highest test concentrations were aligned across all chemicals. For lower concentrations that were not tested in a specific chemical, the barcodes were filled with 0s (indicative of no effects). When being input into the classification methods, the data were truncated to only the highest 5 concentrations such that negative/missing data were not driving the classifications (S7 Fig).

## Supplemental Materials

S1-S10 Figures

S1-S23 Tables

S1 Video: Example planarian behaviors showing the normal gliding behavior compared to two examples of hyperactive behavior.

Supplemental File 1: 95th percentile confidence intervals of the benchmark concentration (in logM) for each planarian endpoint.

## Acknowledgements

The authors thank Dr. Siqi Zhang for helping with data analysis. The authors thank Dr. Michael Lawless (Simulations Plus) for helpful discussions about the use of ANNE and SVMs models and their statistical evaluation and Prof. Øyvind Hammer (University of Oslo, Norway) for helpful discussions of PCoA-LDA.

